# The ‘sexual selection hypothesis’ for the origin of aposematism

**DOI:** 10.1101/2024.09.26.615118

**Authors:** Ludovic Maisonneuve, Thomas G. Aubier

## Abstract

The evolution of aposematism, in which prey exhibit conspicuous signals indicating the presence of anti-predator defenses, is puzzling. Before predators learn to associate the signal with defense, increased visibility makes the conspicuous prey highly vulnerable to predation. Although several hypotheses have been proposed to explain the evolution of aposematism, they often assume that these signals can only be recognized by predators. Yet, many studies show that aposematic signals can also be involved in mate choice. Here, we demonstrate that some aposematic signals may have originally evolved as mating signals driven by sexual selection. In this study, we analyze a mathematical model to explore how sexual selection can drive the evolution of aposematism. We thereby identify key features of this ‘sexual selection hypothesis’ for the origin of aposematism to be tested with empirical data. Our results show that the evolution of conspicuous signals through sexual selection increases the visibility of prey to predators and thus predation pressure. This, in turn, promotes the evolution of defense mechanisms, ultimately leading to aposematism when predators learn to associate the signal with defense. Additionally, we show that when sexual selection drives the evolution of aposematism, it often results in sexual dimorphism in both signaling and defense traits.

## 1 Introduction

Many organisms display conspicuous ‘warning’ signals that warn predators of the presence of a defense mechanism, such as toxins or spines. This anti-predator strategy, known as aposematism, is taxonomically widespread, as it has been documented in plants (Lev-Yadun, 2009, 2024), molluscs (Cortesi and Cheney, 2010), insects (Lindstedt et al., 2017), reptiles (Toledo et al., 2011; Maan and Cummings, 2012), birds (Hedley and Caro, 2022) and mammals (Stankowich et al., 2011; Howell et al., 2021). The initial evolution of aposematism has been widely debated, as it may seem paradoxical at first glance, for at least two reasons. First, a new conspicuous signal evolving from a cryptic trait should be counter-selected because it increases the risk of individuals being detected by predators. Second, a new conspicuous signal is initially rare in the population and offers little protection to the prey since predators are unlikely to have already learned to recognize it as a warning signal. Despite this apparent paradox, many theories have shown that a mutant displaying a warning signal can be favored in a population of cryptic prey, thus explaining the initial evolution of aposematism. Notably, all the underlying mechanisms, detailed below, are based on the assumption that prey are initially defended. This is likely to be the case when defenses increase the prey’s chances of surviving predator attacks (Leimar et al., 1986), with the resulting benefit outweighing the costs of maintaining these defenses.

The initial evolution of aposematism may be promoted by kin selection (Fisher, 1930; Benson, 1971; Waldman and Adler, 1979; Gittleman and Harvey, 1980; Harvey et al., 1982; Leimar et al., 1986; Broom et al., 2006; Scaramangas and Broom, 2022; Scaramangas et al., 2023). When a defended individual displaying a warning signal is attacked by a predator, the predator may learn to avoid the signal. This learning process reduces predation on the individual’s relatives, provided they are likely to be nearby and display the same warning signal. Therefore, for kin selection to favor the initial evolution of aposematism, a predator attacking a prey individual must be likely to encounter the individual’s relatives. This is the case for a fine-scale population structure, such as that caused by family grouping or low dispersal (Mallet and Singer, 2008).

Other theories have shown that the initial evolution of aposematism may be based on individual selection rather than kin selection. A defended individual may benefit from displaying a warning signal if it survives a first attack by a predator, leading the predator to avoid attacking it in future encounters (Järvi et al., 1981; Wiklund and Järvi, 1982; Sillén-Tullberg and Bryant, 1983; Sillén-Tullberg, 1985; Tullrot, 1994). A defended individual displaying a warning signal may also have higher fitness than cryptic individuals simply because of the protection provided by aposematism (Servedio, 2000; Guilford, 1985). This advantage can be further enhanced when defended prey are located alongside undefended prey, as the warning signal may distinguish defended prey from undefended prey (Sherratt and Franks, 2005; Franks et al., 2009). This protection provided by aposematism occurs when predators have learned to avoid the warning signal by attacking other defended prey displaying the signal. Since this process relies on the warning signal being sufficiently common in the prey population, it cannot explain the evolution of aposematism from a single mutant displaying a warning signal (Servedio, 2000). Nonetheless, a new warning signal could nevertheless become sufficiently common to be favored because of stochasticity (Mallet and Singer, 2008) or other aspects of predator psychology, such as dietary conservatism or neophobia (Lee et al., 2010; Aubier and Sherratt, 2015). Furthermore, the rise in frequency of a new warning signal may also be facilitated by innate biases in predator learning, particularly when predators can easily recognize and remember conspicuous patterns (e.g. Halpin et al. (2008a,b)). It should be noted that the different mechanisms explaining the emergence of aposematism are not mutually exclusive and may operate in combination.

Most of the mechanisms explaining the initial evolution of aposematism rely on the assumption that warning signals are only recognized by predators. However, warning signals can also serve as the basis for mate choice (Rojas et al., 2018), as shown by numerous empirical studies (e.g., Reynolds and Fitzpatrick, 2007; Maan and Cummings, 2008, 2009; Nokelainen et al., 2012; Finkbeiner et al., 2014b; Jiggins et al., 2001; Estrada and Jiggins, 2008; Gordon et al., 2015; Chouteau et al., 2017). Such mate choice can generate sexual selection, i.e., selection through differential mating success, that is known to have favored the evolution of colorful signals (Hill, 1991; Stuart–Fox and Ord, 2004; Olsson et al., 2013; Maan and Sefc, 2013; Dale et al., 2015). It has therefore been proposed that some warning signals may have evolved initially as sexual signals (Mallet and Singer, 2008; Ruxton et al., 2004). According to this scenario, sexual selection may eventually lead to aposematism: when prey are defended, predators may associate these sexual signals favored by sexual selection with the prey’s defenses to avoid attacking them.

While many mathematical models have studied the effects of sexual selection on the diversity of warning signals (Tazzyman and Iwasa, 2010; Boussens-Dumon and Llaurens, 2021; Ponkshe and Endler, 2022; Yeager and Penacchio, 2023; Maisonneuve et al., 2021, 2023), none has investigated the role of sexual selection in the initial evolution of aposematism. Yet, sexual selection favoring signal evolution should impact the evolution of aposematism by affecting the co-evolutionary dynamics between defense and signal (as in Leimar et al., 1986; Broom et al., 2008, in a case without sexual selection). Indeed, the evolution of a conspicuous sexual signal increases predation pressure, which should favor defense evolution. In turn, defense evolution increases the efficiency of predator learning, which should reduce predation pressure and thus affect signal and defense evolution.

In this study, we develop a mathematical model to investigate the coevolution of signal and defense, and aim to examine the impact of sexual selection on the evolution of aposematism. Our results indicate that sexual selection can favor the initial evolution of aposematism, even when prey originally had no defense against predators. Assuming that the female is the choosy sex, we also show that the evolution of aposematism through sexual selection is likely to result in strong sexual dimorphism in signal and defense, with a more pronounced effect on defense.

## 2 Model

We consider a sexually reproducing haploid population. Each individual expresses a trait *s*_•_, which determines the intensity of a signal it displays (note that, here and hereafter, the symbol • refers to the trait of a focal individual) (main symbols are listed in Table 1). Such a signal may take the form of a colorful pattern or exaggerated ornamentation. If an individual does not display a signal, then *s*_•_ = 0. We assume that the signal can be recognized by both predators and mate-seeking females. Each individual also expresses a trait *d*_•_, which determines the individual’s defense level, i.e., the efficiency of the defense mechanism in increasing the prey’s survival probability in the event of a predator attack. For example, individuals may be poisonous, have an unpalatable taste, or may protect themselves by stinging. If an individual is completely harmless to predators, then *d*_•_ = 0. We also assume that predators decide whether to attack prey based on their previous experiences, and that they may learn to associate the signal with defense, so that they avoid attacking prey displaying the signal.

**Table 1:** Key symbols and their definitions.

| Model parameters |  |
| --- | --- |
| $\gamma_{cs1}, \gamma_{cs2}$ | Strength of the linear and quadratic production costs associated with the signal |
| $\gamma_{cd1}, \gamma_{cd2}$ | Strength of the linear and quadratic production costs associated with the defense |
| $p$ | Encounter rate between prey and predators |
| $P_{D0}$ | Probability that a predator will detect a prey without signal |
| $\beta_D$ | Extent to which the signal intensity increases the probability of the prey being detected |
| $\beta_N$ | Extent to which the signal intensity increases the probability of the signal being noticed |
| $\beta_E$ | Extent to which the defense level increases the probability of prey escape |
| $\beta_L$ | Extent to which the defense level increases predator avoidance learning |
| $\lambda$ | Predator forgetting rate |
| $\rho$ | Strength of sexual selection |
| $N$ | Population size |
| Variables describing traits in the model with sex-independent traits |  |
| $\mathbf{x} = (s, d)$ | Resident trait values for signal and defense |
| $\mathbf{x}_\bullet = (s_\bullet, d_\bullet)$ | Mutant/focal individual trait values for signal and defense |
| Variables describing traits in the model with sex-dependent traits |  |
| $\mathbf{x} = (s_m, d_m, s_f, d_f)$ | Resident trait values for male and female signal and defense |
| $\mathbf{x}_\bullet = (s_{m\bullet}, d_{m\bullet}, s_{f\bullet}, d_{f\bullet})$ | Mutant/focal individual trait values for male and female signal and defense |

We consider a Wright-Fisher model with a constant population size *N* that we assume to be large enough to ignore stochastic effects. We assume the following life cycle: (1) over a standardized time interval of 1, individuals survive or die depending on predation and the intrinsic costs associated with their traits; (2) at the end of the time interval, females choose which males to mate with based on their signal; (3) mating pairs produce a fixed number of offspring; and (4) adults die so that generations do not overlap.

We investigate whether the combined action of sexual selection and viability selection can favor the evolution of signal and defense in a population, leading to predators to avoid attacking prey displaying the signal. In the following, we first describe how individual traits (*s*_•_ and *d*_•_) affect different components of individual fitness, including production costs, predation and sexual selection (sections 2.1 to 2.4). We then describe how these components together determine the fitness associated with a phenotype, depending on whether traits are sex-independent or sex-dependent (section 2.5). Finally, we explain how we analyze the co-evolutionary dynamic in the model (section 2.6).

### 2.1 Production costs

Individuals allocate a part of their metabolism to the production of signal and defense. This allocation may occur at the expense of other biological functions (*e*.*g*., tissue growth, thermoregulation, antioxidant defense). In line with empirical evidence (*e*.*g*., Bryant and Julkunentiitto, 1995; Grill, 1999; Rigby and Jokela, 2000; Zalucki et al., 2001; Marak et al., 2003; Ojala et al., 2007; Lindstedt et al., 2010; Blount et al., 2012, 2023), we therefore assume that the production of the signal or defense induces constitutive costs. In our model, these costs raise the mortality rate of an individual with traits *s*_•_ and *d*_•_ by

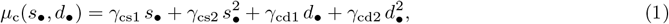

which increases as the individual’s traits *s*_•_ and *d*_•_ increase. The parameters *γ*_cs1_ and *γ*_cs2_ (resp. *γ*_cd1_ and *γ*_cd2_) tune the strength of the linear and quadratic production costs associated with the signal (resp. defense).

### 2.2 Predation

#### 2.2.1 Prey detection by the predator

We assume that prey encounter predators at a constant rate *p*. Upon encounter, the predator can detect the prey, depending on the intensity *s*_•_ of the prey’s signal. The detection probability *P*_D_ increases with the individual’s signal intensity *s*_•_ and is given by

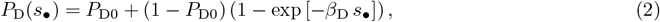

where *P*_D0_ is the probability that a predator will detect a prey without signal (i.e., a prey with trait *s*_•_ = 0) and *β*_D_ quantifies the extent to which the signal intensity increases the probability of detection.

#### 2.2.2 Attack of the prey by the predator

If a predator detects a prey, it chooses whether or not to attack. A predator chooses to attack based on the prey’s signal and its own previous experiences. We assume that the predator notices the prey’s signal with probability

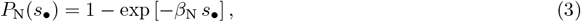

which increases with the intensity *s*_•_ of the prey’s signal. The parameter *β*_N_ quantifies how much the signal intensity increases the probability of the signal being noticed. If the predator does not notice the signal (with probability 1 − *P*_N_(*s*_•_)), it always attacks the prey. By contrast, if the predator notices the signal (with probability *P*_N_(*s*_•_)), it will only attack the prey if it has not learned to associate the prey’s signal with defense, i.e., if it does not yet consider the prey’s signal to be a warning signal.

The resulting attack probability on a prey with a signal intensity *s*_•_ is then given by

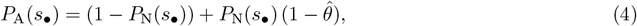

where 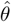 denotes the proportion of experienced predators that have learned to associate the prey’s signal with defense and thus avoid attacking signaling prey. To derive the expression for the proportion of experienced predators 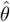, we make assumptions about predator avoidance learning, as explained in the next subsection.

#### 2.2.3 Predator avoidance learning

Following Ferreira and Marcon (2014), we assume that predators learn to avoid prey displaying the signal but also forget at rate *λ*. We assume that after attacking prey for which the predator has noticed the signal, the predator learns to avoid prey displaying the signal with probability *P*_L_(*d*_•_). In line with empirical evidence (*e*.*g*., Lindström et al., 1997; Skelhorn and Rowe, 2006b), the avoidance learning probability *P*_L_(*d*_•_) increases with the defense level *d*_•_ of the prey, so that

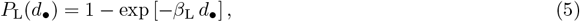

where *β*_L_ modulates the effect of the prey defense level *d*_•_ on predator avoidance learning. Note that after attacking a defenseless prey (i.e., a prey with *d*_•_ = 0), the predator has no negative experience and does not learn to avoid signaling prey (*P*_L_(0) = 0).

We assume that the prey population is much larger than the predator population, and that the proportion of experienced predators reaches its equilibrium value before predation significantly changes the trait distribution in the prey population. Under these assumptions, the proportion of experienced predators at equilibrium is given by

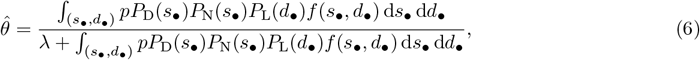

where *f* is the trait distribution in the prey population (see Appendix A1 for more details on the analytical derivations).

Eq. (6) shows that a balance between learning and forgetting determines the equilibrium proportion of experienced predators 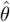. Indeed, the proportion of experienced predators 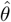 decreases with the forgetting rate *λ*. In contrast, the proportion of experienced predators 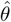 is high when predators attack prey and notice their signal, and when they efficiently learn to associate the prey signal with defense. Overall, 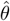 depends on the prey’s signal and defense. In other words, signal and defense within the prey population together determine predator avoidance learning, as shown empirically (Halpin et al., 2008a,b).

#### 2.2.4 Prey escape

As detailed above, predators attack the prey they encounter with probability *P*_A_(*s*_•_). We assume that the prey can escape an attack with a probability *P*_E_(*d*_•_) that increases with the prey’s defense level *d*_•_ (in line with empirical studies; Wiklund and Järvi, 1982; Skelhorn and Rowe, 2006a,c) so that

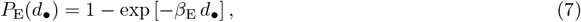

where *β*_E_ quantifies how the defense level increases the probability of prey escape. We assume that if the prey does not succeed in escaping, it gets killed by the predator.

#### 2.2.5 Overall predation pressure

Overall, predation raises the mortality rate of an individual with traits *s*_•_ and *d*_•_ by

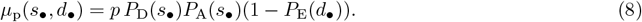

### 2.3 Probability of surviving to reproductive maturity

At the end of a standardized time interval of 1, survivors gather to mate. The survival probability of an individual with traits *s*_•_ and *d*_•_ is

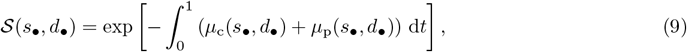

where *t* is the time within the generation.

By substituting the expression of *μ*_c_(*s*_•_, *d*_•_) and *μ*_p_(*s*_•_, *d*_•_) from eqs. (1) and (8) into Eq. (9) we obtain

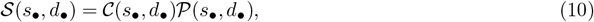

where

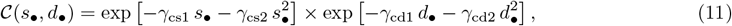

and

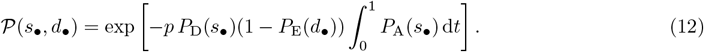

Note that in the general case, 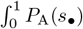 cannot be calculated analytically because *P*_A_(*s*_•_) changes over time. This variation occurs because *P*_A_(*s*_•_) depends on the proportion of experienced predators 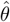, which itself is influenced by the distribution of traits. The distribution of traits shifts over the course of a generation as individuals with different traits experience varying death rates. To ensure the model remains mathematically tractable, we assume that the distribution of traits remains nearly constant during a generation (and therefore that the sex ratio remains balanced) allowing us to write 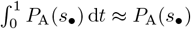 leading to:

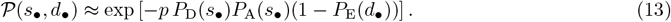

This assumption is relaxed in individual-based simulations and our results remain qualitatively unchanged (see Appendix A2 for details).

### 2.4 Sexual selection

The surviving adults come together to mate. We consider a panmictic population, where each individual encounters others with equal probability. We assume that all surviving females mate equally and produce the same number of offspring. We also assume that females are the choosy sex and have the same mating preference, so that they prefer males with the most intense signal. The mating probability of a male with signal intensity *s*_•_ is therefore proportional to its attractiveness

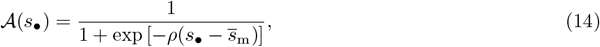

where *ρ* describes the strength of female preference for more intense signal, and 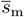 is the mean male signal in the population. The attractiveness of a focal male, and therefore his mating probability, increases as the intensity of his signal exceeds the population mean.

### 2.5 Fitness

#### 2.5.1 Individual fitness

The reproductive output *f*_m_ and *f*_f_ of males and females with traits *s*_•_ and *d*_•_ are then given by

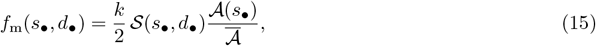

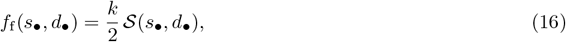

where *k >* 0 is the number of offspring produced by each female, assumed to be constant, and is such that the average reproductive output of both males and females in the population exceeds one. The term 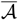 is the average attractiveness of males, such that 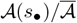 gives the probability that a male with signal *s*_•_ successfully mates.

The fitnesses *w*_m_ and *w*_f_ of males and females with traits *s*_•_ and *d*_•_ are given by

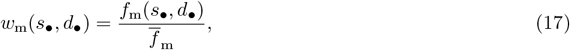

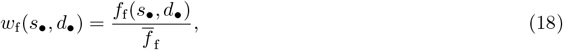

where 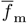 and 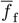 are the mean fertilities of males and females in the population, respectively. Due to normalization, 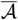 and *k* introduced in equations (15) and (16) do not affect fitnesses.

#### 2.5.2 Phenotypic fitness

The evolutionary dynamics hinge on the fitness of individuals, characterized by their signaling and defense traits, and therefore depend on whether the traits expressed in females and males are encoded by the same genes or by different genes. We thus consider two versions of the model that differ in how we define the phenotypic space: a model with sex-independent traits where the population is necessarily sexually monomorphic, and a model with sex-dependent traits where the population can be sexually dimorphic.

##### Model with sex-independent traits

In this model, the signal and the defense of an individual does not depend on whether the individual is a male or a female. Each individual is characterized by a phenotypic vector **x**_•_ = (*s*_•_, *d*_•_). Since the sex ratio is assumed to be balanced, the fitness associated with phenotype **x**_•_ is given by:

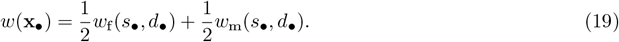

##### Model with sex-dependent traits

In this model, the signal and the defense of an individual depends on whether the individual is a male or a female. Each individual is characterized a phenotypic vector **x**_•_ = (*s*_f•_, *d*_f•_, *s*_m•_, *d*_m•_). If the individual displaying phenotype **x**_•_ is female (resp. male), it expresses the traits *s*_f•_ and *d*_f•_ (resp. *s*_m•_ and *d*_m•_). The fitness associated with the phenotype **x**_•_ is then given by:

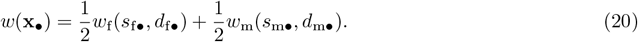

### 2.6 Model analyses

#### 2.6.1 Invasion analyses

We assume that mutations are rare and have little effect on phenotype. Based on this assumption, the phenotypic evolutionary dynamic can be inferred from the invasion of rare mutants into a resident population (Champagnat et al., 2006; Mullon and Lehmann, 2019).

We denote *w*_inv_(**x**_•_, **x**) the invasion fitness of a mutant, i.e., the fitness of a rare mutant with phenotype **x**_•_ in a population with phenotype **x**. The invasion fitness of a mutant with phenotype **x**_•_ in a resident population with phenotype **x** (with **x** = (*s, d*) or **x** = (*s*_f_, *d*_f_, *s*_m_, *d*_m_) depending on whether we consider sex-independent or sex-dependent traits) is given by

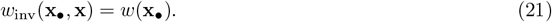

Note that *w*(**x**_•_), defined in equations (19) and (20), depends on the resident phenotype **x** because the proportion of experienced predators 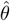 actually depends on the resident phenotype **x** in the prey population (see Appendix A3).

We assume that mutations independently affect each trait. The change in phenotypic traits per unit of time can then be approximated by

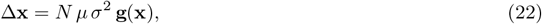

where *μ* is the mutation rate, *σ*^2^ is the variance in mutational effects and **g**(**x**) is the selection gradient (Dieckmann and Law, 1996; Champagnat et al., 2006; Metz, 2011; Ripa and Dieckmann, 2013). The selection gradient is defined by

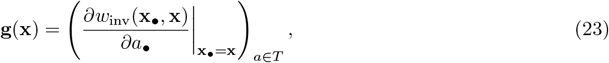

where *T* = {*s, d*} or *T* = {*s*_m_, *s*_f_, *d*_m_, *d*_f_} depending on whether we consider sex-independent or sex-dependent traits. The selection gradient describes the direction of evolutionary changes within the phenotypic space. The expressions of the selection gradients in models with sex-independent or sex-dependent traits are shown in Appendix A4.

#### 2.6.2 Assumptions on the ancestral population

We consider that the ancestral population has reached a state of evolutionary equilibrium, with sexual selection playing no role. In the absence of sexual selection, we assume that signaling is deleterious and that the ancestral population is therefore not aposematic (see Appendix A6). Sexual selection on the signal then occurs, disrupting this equilibrium and affecting the evolution of signaling and defense.

In a first analysis, we assume that the ancestral population is not defended at its evolutionary equilibrium, that is, the selection gradients for defenses within this ancestral population are negative 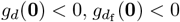 and 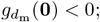 see eqs. (A8), (A10) and (A12), and note that this occurs under the same conditions for both sex-independent and sex-dependent traits). This means that, ancestrally, selection inhibits the evolution of defense, which is a condition that prevents the evolution of aposematism. In this ‘worst-case scenario’, we investigate whether sexual selection can enable the initial evolution of aposematism through the evolution of signaling and defense in the prey. We analytically determine the conditions required for the emergence of signal and defense from the expression of the selection gradients. We then numerically estimate the phenotype **x**^∗^ reached at evolutionary equilibrium when sexual selection takes place. At this evolutionary equilibrium, we assess the degree of aposematism as the proportion of experienced predators 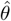 that avoid attacking prey displaying the signal.

In a second analysis, we take a more general approach and consider that defense may evolve in the ancestral population even without sexual selection. As in the previous analysis, we numerically estimate the phenotype reached at evolutionary equilibrium. But this time, we choose parameter values at random (see details of the procedure in Appendix A5) and assess whether sexual selection correlates with the evolution of signal and defense (i.e., with aposematism).

## 3 Results

### 3.1 Evolution of aposematism in ancestrally defenseless populations

#### 3.1.1 Sexual selection and predation jointly promote the evolution of aposematism

We consider the model with sex-independent traits, in which the population is assumed to be sexually monomorphic. As explained above, we assume that when there is no sexual selection on the signal, prey defense is counter-selected; therefore, the population is ancestrally defenseless. We show that the condition for the initial evolution of the signal (that is, in a population characterized by *s* = 0 and *d* = 0) is then

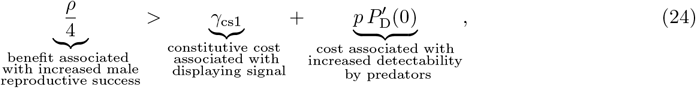

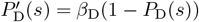. See Appendix A7.1 for details on the derivation.

Eq. (24) shows that the costs inhibiting signal evolution are twofold: displaying the signal comes with constitutive costs and increases detectability by predators. These costs may prevent the initial evolution of the signal in the population (*e.g*., as in Fig. 1a). Nonetheless, Eq. (24) also reveals that the signal can be favored by sexual selection if females sufficiently prefer it. Indeed, a high reproductive success of signaling males (high *ρ*) can offset the costs associated with signaling, favoring signal evolution (*e.g*., as in Fig. 1b-c).

**Figure 1:**
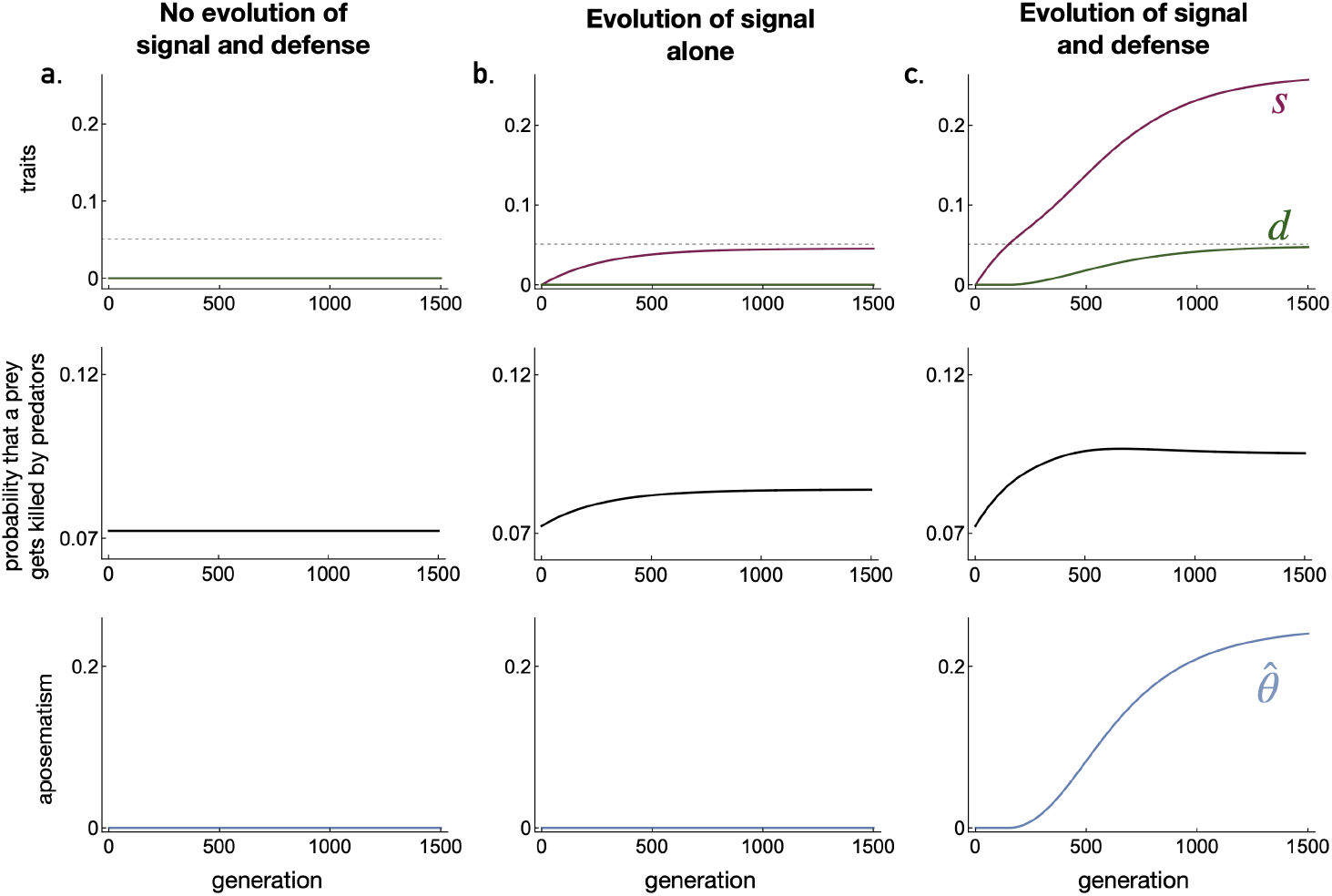
The evolution of signal and defense in the model with sex-independent traits. Temporal dynamics under three different scenarios of mean signal intensity (*s*) and defense level (*d*), the probability that a prey gets killed by predators, and aposematism quantified as the proportion of experienced predators. **a**. When females express weak preference for male signal (*ρ* = 1.4), costs associated with signal prevent its evolution. **b**. When females express moderate preference for male signal (*ρ* = 1.59), sexual selection promotes the evolution of signal despite the associated costs. Nevertheless, the signal does not become intense and does not increase predation pressure enough for defense to be favored. **c**. When females express strong preference for male signal (*ρ* = 1.7), sexual selection promotes the evolution of an intense signal. The evolution of an intense signal increases predation pressure and promotes the evolution of defense against predators. In all panels, the dashed line represents the threshold value outlined in Eq. (26); defense can initially evolve when the intensity of the signal in the population has reached this threshold value (as occurs in panel **c**., but not in panels **a**.-**b**. even if we run the simulation for longer). Parameter values are: *γ*_cs1_ = 0.1, *γ*_cs2_ = 0.35, *γ*_cd1_ = 0.08, *γ*_cd2_ = 0.1, *β*_D_ = 1.25, *β*_N_ = 17.5, *β*_E_ = 0.9, *β*_L_ = 0.5, *λ* = 0.01, *P*_D0_ = 0.25, *p* = 0.3.

The evolution of the signal is associated with an increase in the probability of prey being killed by predators (via an increase in detectability). This affects selection on prey defense. Indeed, the condition for the initial evolution of defense once the signal has evolved (that is, in a population characterized by *s >* 0 and *d* = 0) is

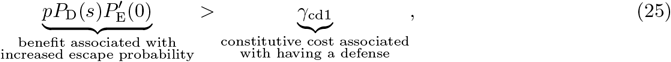

where 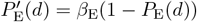. See Appendix A7.2 for details on the derivation.

Eq. (25) shows that defense can evolve despite its constitutive cost because it enables prey to escape predators. Defense is particularly beneficial when the predation pressure is high (high *p P*_D_(*s*)), including when prey are highly detectable (high *P*_D_(*s*)). As the evolution of the signal through sexual selection increases prey detectability and therefore predation pressure, it favors the initial evolution of defense. We show that the intensity of the signal must exceed a threshold value 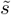 for predation pressure to be high enough to favor defense evolution (see Appendix A7.1 for derivation):

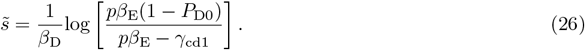

When signal intensity remains below this threshold, there is no defense evolution (*e.g*., as in Fig. 1a-b). In contrast, when signal intensity exceeds this threshold, predation pressure becomes sufficiently high to favor the initial evolution of defense (*e.g*., as in Fig. 1c).

As signaling and defense evolve, predators may learn to avoid prey that display the signal; the prey’s signal is considered as a warning signal by experienced predators, and aposematism has thus evolved (*e.g*., as in Fig. 1c). The evolution of prey defense, in turn, influences the selective pressures acting on the signal. To investigate this feedback on signal evolution, let us consider the selection gradient on the signal in a population characterized by *s >* 0 and *d >* 0:

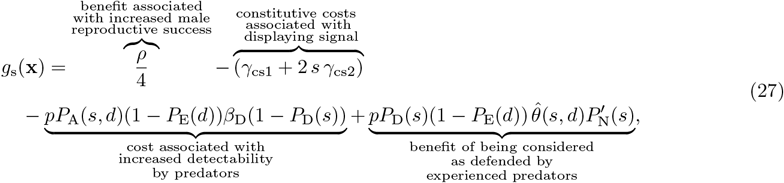

where 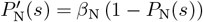 (see Eq. (A7)). The selection gradient *g*_s_(**x**) indicates the direction of selection on signal intensity. For instance, a more intense signal is favored when this selection gradient is positive.

The first three terms of Eq. (27) are analogous to the terms found in Eq. (24) describing the condition for the initial evolution of signaling. In contrast, the last term of Eq. (27) is equal to zero when *d* = 0 and is therefore not found in Eq. (24). It shows that once signaling and defense have both evolved, aposematism promotes the evolution of a more intense signal. This is because signal intensity increases the chance of the signal being noticed and avoided by experienced predators that consider the signal as a warning signal. As signal intensity increases, aposematism strengthens, reducing the probability of prey being killed by predators (*e.g*., as in Fig. 1c). This benefit is high when most predators are experienced and avoid attacking prey that display the signal (high 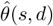), and when signal intensity greatly increases the probability of predators noticing the signal (high 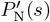). The proportion 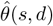 of experienced predators that consider the signal as a warning signal is given by

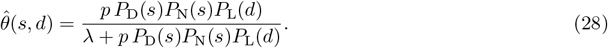

See Appendix A3 for details on the derivation. The proportion of experienced predators is high when predators frequently attack prey and notice their signal (high *p P*_D_(*s*)*P*_N_(*s*)), when they efficiently learn to associate the signal with defense (high *P*_L_(*d*)), and when they have a low forgetting rate (low *λ*).

To further study the impact of aposematism on predation pressure, we investigate the effects of predator forgetting rate *λ* on traits and the probability of prey being killed by predators at evolutionary equilibrium. When the predator forgetting rate is low, the proportion of experienced predators becomes high (Fig. 2), corresponding to a case with strong aposematism. As described earlier, we find that such strong aposematism leads to a low probability of prey being killed by predators (Fig. 2). However, sensitivity analyses indicate that, on average, the predator forgetting rate has a very weak effect on the probability of a prey being killed by a predator (see Tab. A1). This is because protection provided by aposematism favors the evolution of a signal with high intensity, which increases predation pressure, partially offsetting the benefits of aposematism.

**Figure 2:**
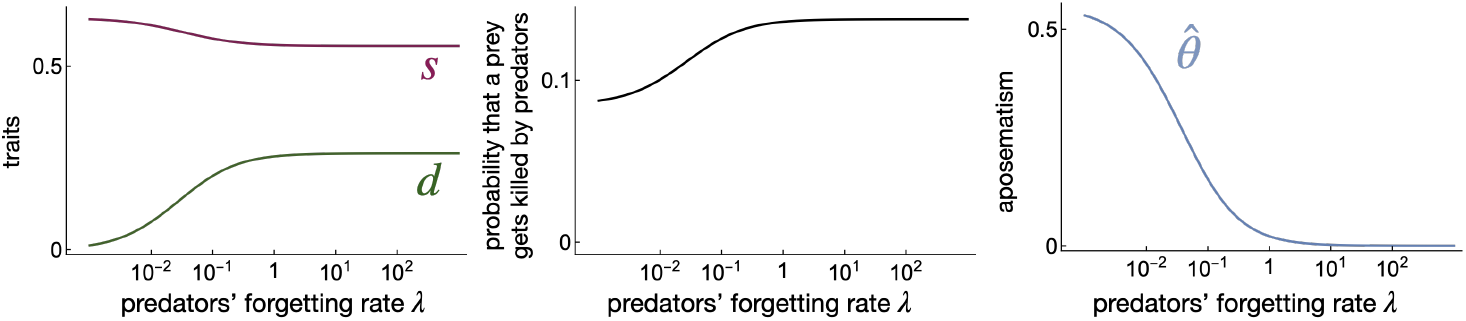
Impact of predators’ forgetting rate on the evolution of signal and defense in the model with sex-independent traits. Mean signal intensity *s* and defense level *d*, probability that a prey gets killed by predators, and aposematism quantified as the proportion of experienced predators, depending on the predators’ forgetting rate *λ*. Parameter values are: *ρ* = 2.4, *γ*_cs1_ = 0.1, *γ*_cs2_ = 0.35, *γ*_cd1_ = 0.08, *γ*_cd2_ = 0.1, *β*_D_ = 1.25, *β*_N_ = 17.5, *β*_E_ = 0.9, *β*_L_ = 0.5, *P*_D0_ = 0.25, *p* = 0.3.

#### 3.1.2 The evolution of aposematism results in sexual dimorphism in signaling and defense

In the previous section, we demonstrated that the interplay between sexual selection and predation can favor the evolution of aposematism. However, we assumed that males and females necessarily express the same signal intensity and defense level, so that sexual monomorphism is maintained in the population. Since sexual selection only affects the reproductive success of males in our model, we now investigate how aposematism can evolve in a population that may be sexually dimorphic. We consider an extreme case where males and females express different signaling and defense traits that are encoded by different genes. In this context, the female signal is not under sexual selection.

We show that the condition for the initial evolution of signal and defense in males (that is, in a population characterized by *s*_f_ = 0, *d*_f_ = 0, *s*_m_ = 0 and *d*_m_ = 0, and then by *s*_f_ = 0, *d*_f_ = 0, *s*_m_ *>* 0 and *d*_m_ = 0) is qualitatively similar to that for the initial evolution of the signal and defense in the previous model with sex-independent traits (see Appendix A8.1).

Once signal and defense have evolved in males, the condition for the initial evolution of female signal (that is, in a population characterized by *s*_f_ = 0, *d*_f_ = 0, *s*_m_ *>* 0 and *d*_m_ *>* 0) is

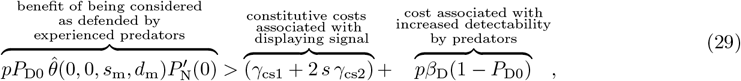

where 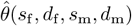 is the proportion of experienced predators when the prey population is characterized by the phenotypic vector (*s*_f_, *d*_f_, *s*_m_, *d*_m_). Since we assume that the distribution of traits remains nearly constant throughout a generation for the sake of analytical tractability, we assume that the sex ratio remains balanced over time (this hypothesis is relaxed in the individual-based model presented in Appendix A2). Consequently, we have

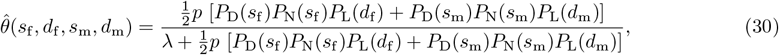

See Appendix A3 for details on the derivation.

Eq. (29) indicates that despite the constitutive costs and the increased predation risk associated with increased detectability, female signaling can be beneficial when there is aposematism mediated by signaling and defense in males. Here, aposematism allows signaling females to be identified as defended by predators, even if they are defenseless. If this benefit is substantial, female signal can evolve (*e.g*., as in Fig. 3). This is the case, for instance, when predators have a low forgetting rate (low *λ*) and male defense enhances predator avoidance learning (high *β*_L_) leading to a high proportion of experienced predators (Figs. 4 and A1a). Furthermore, the advantage of being identified as defended by predators increases with the effect of the signal on the probability of it being detected *β*_N_, promoting the evolution of more intense female signal (Fig. A1b). Interestingly, we observe a high degree of sexual dimorphism in signal at evolutionary equilibrium (Fig. 4).

**Figure 3:**
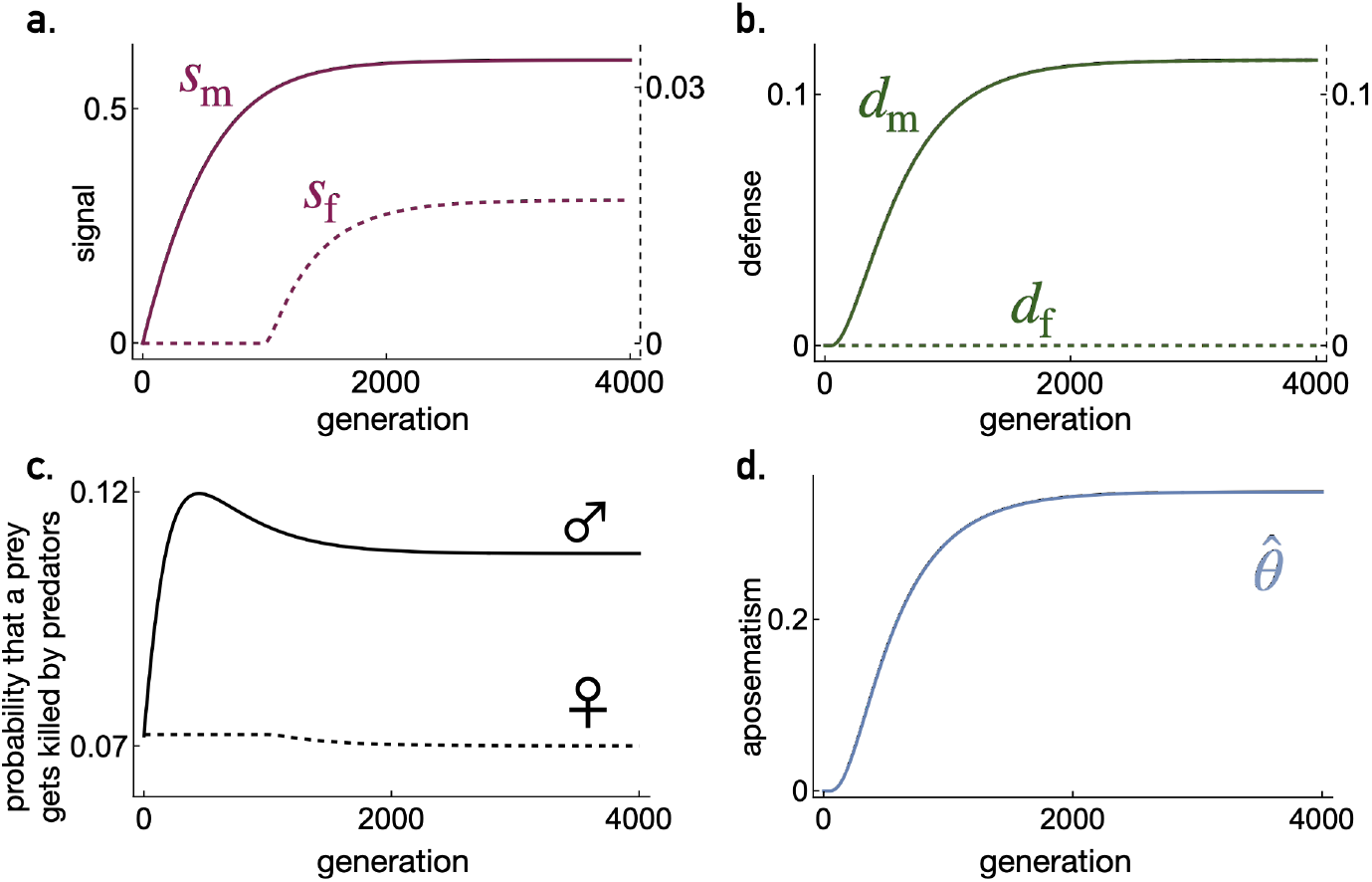
The evolution of signal and defense in males and females in the model with sex-dependent traits. Temporal dynamics of **a**. mean male and female signal intensity, **b**. mean male and female defense level, **c**. the probability that a male or female prey gets killed by predators, and **d**. aposematism quantified as the proportion of experienced predators. Parameter values are: *ρ* = 1.2, *γ*_cs1_ = 0.1, *γ*_cs2_ = 0.35, *γ*_cd1_ = 0.08, *γ*_cd2_ = 0.1, *β*_D_ = 1.25, *β*_N_ = 17.5, *β*_E_ = 0.9, *β*_L_ = 0.5, *λ* = 0.01, *P*_D0_ = 0.25, *p* = 0.3.

**Figure 4:**
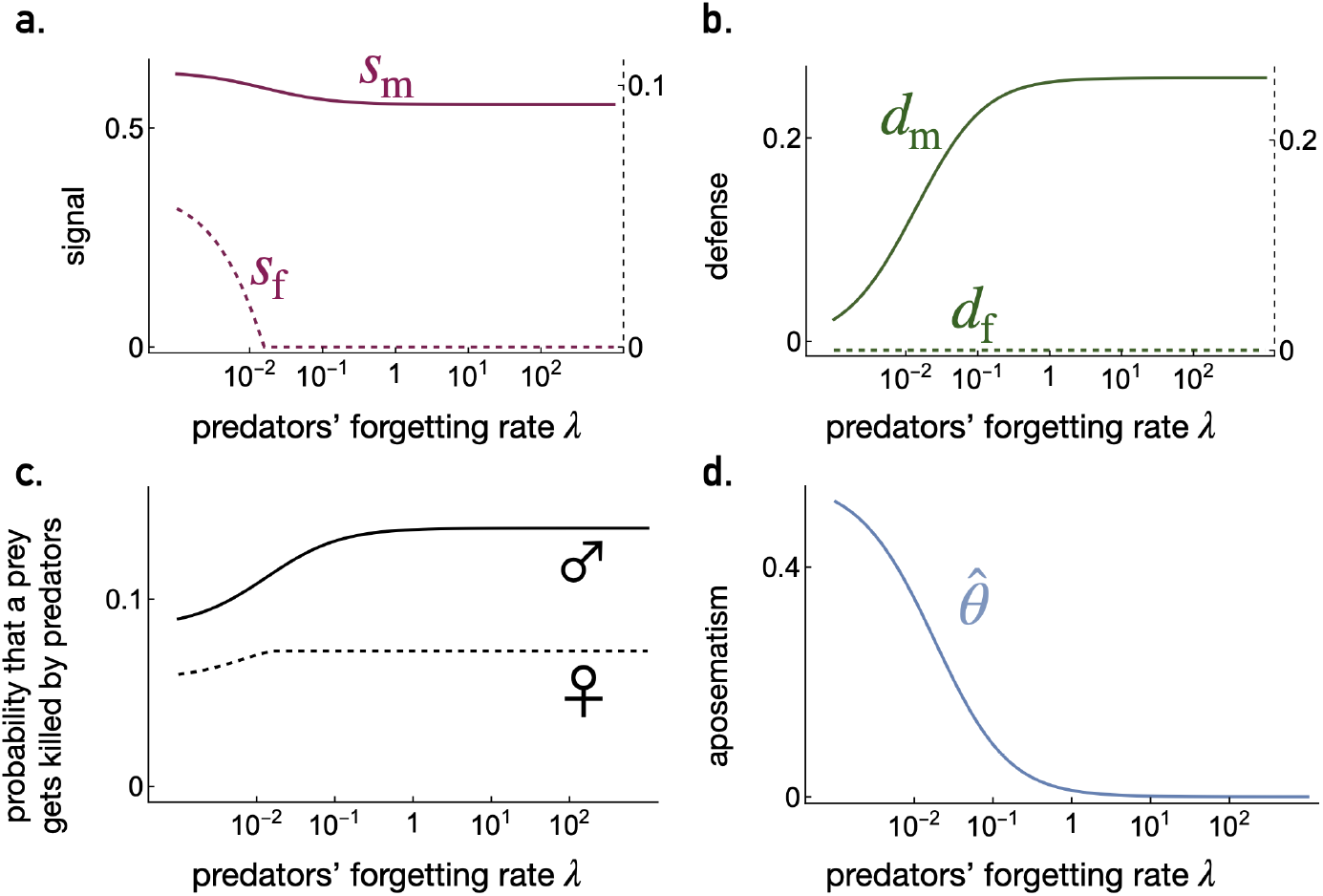
Impact of predators’ forgetting rate on the evolution of signal and defense in males and females in the model with sex-dependent traits. **a**. Mean male and female signal intensity, **b**. mean male and female defense level, **c**. the probabilities that a male and a female get killed by predators, and **d**. aposematism quantified as the proportion of experienced predators, depending on the predators’ forgetting rate *λ*. Parameter values are: *ρ* = 1.2, *γ*_cs1_ = 0.1, *γ*_cs2_ = 0.35, *γ*_cd1_ = 0.08, *γ*_cd2_ = 0.1, *β*_D_ = 1.25, *β*_N_ = 17.5, *β*_E_ = 0.9, *β*_L_ = 0.5, *P*_D0_ = 0.25, *p* = 0.3.

This is due to sexual selection, which results in the intensity of the male signal being greater than that of the female signal.

We next examine whether female defense can evolve once signaling (in both males and females) and male defense have evolved. We show in Appendix A8.3 that female defense cannot evolve if the population is ancestrally defenseless. Remember that we assume that in the absence of signaling, the constitutive cost of defense exceeds the benefit associated with increased escape probability. Yet, the evolution of female signal reduces predation pressure on females by allowing them to benefit from aposematism (Fig. 3). This further diminishes the benefit of female defense (the increase in escape probability), inhibiting its evolution. As a result, the population displays strong sexual dimorphism in defense, with females having no defense at all.

Our findings indicate that the evolution of aposematism through sexual selection on male signal should be associated with sexual dimorphism in both signaling and defense. To assess the reliability of our results, we conducted individual-based simulations allowing the distribution of traits and the sex ratio to vary throughout a generation, and found consistent results (see Fig. A2).

Remember that we consider an extreme scenario, where male and female traits are encoded by entirely separate genes and where sexual selection affects only the evolution of male signal. Sexual dimorphism may therefore be less pronounced in natural populations where these assumptions are unmet. Nonetheless, in populations where sexual selection is at the origin of aposematism, our model predicts sexual dimorphism in both signaling and defense, at least to some extent.

### 3.2 Evolution of aposematism in defended and defenseless populations

We now investigate the evolution of aposematism by sexual selection, without making any restricting assumptions about the ancestral population. We consider the possibility that defense may be favored before sexual selection occurs, so that prey are ancestrally defended. As in the previous analysis, we consider sex-dependent traits, but this time we conduct a sensitivity analysis to assess the impact of the different parameters on aposematism and sexual dimorphism. More specifically, we randomly choose parameter values within a given range (see Appendix A5 for details) and then determine numerically trait values at evolutionary equilibrium. At this equilibrium, we compute the level of aposematism 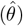 along with measures of sexual dimorphism in both signal and defense (*δ*_s_ and *δ*_d_; see Appendix A9 for details).

Based on 1,000,000 simulations with different combinations of parameter values, we find that strong sexual selection is associated with high levels of aposematism (see Fig. 5a; Pearson correlation coefficient: 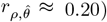. This finding confirms our previous result, which suggests that sexual selection can promote the evolution of aposematism. Furthermore, we find that the evolution of a high level of aposematism is correlated with a high level of sexual dimorphism in both signal and defense (see Fig. 5b and 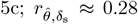 and 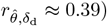. Note that the correlation between the level of aposematism and the level of sexual dimorphism in defense is particularly strong, as expected given our previous model analysis. Altogether, this reinforces our previous findings that aposematism evolving through sexual selection should be associated with sexual dimorphism in both signal and defense.

**Figure 5:**
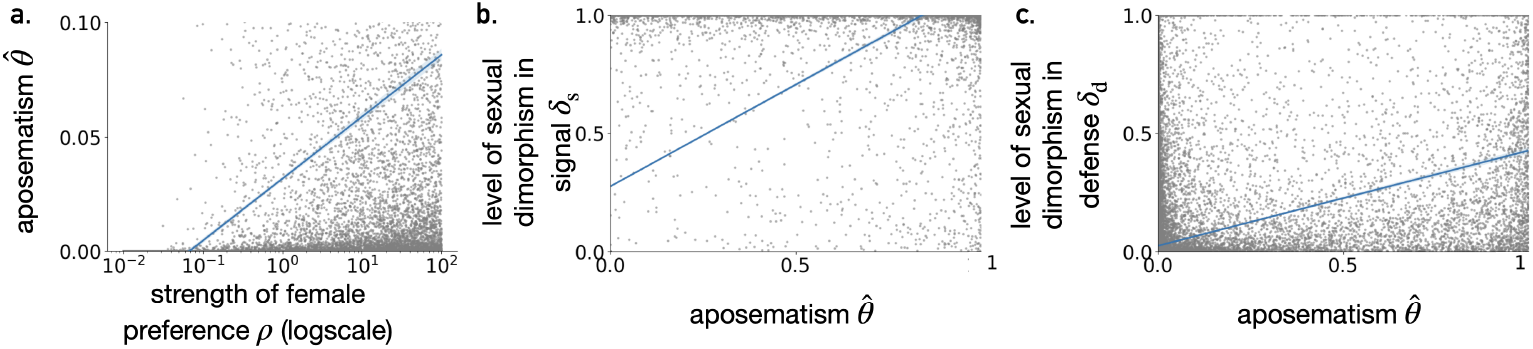
The impact of female preference on the evolution of aposematism and its role in the evolution of sexual dimorphism. Each dot represents the outcome of a numerical simulation at evolutionary equilibrium. Simulations were performed using a randomly selected combination of parameter values according to the procedure described in section A5. In particular, values of log[*ρ*] are randomly sampled following a uniform distribution between −2 and 2, ensuring that the parameter values span multiple orders of magnitude. We represent: **a**. the proportion of experimented predators 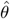 depending on the strength of female preference *ρ*, **b**. 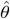 depending on the level of sexual dimorphism in signal *δ*_s_ and **c**. 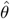 depending on the level of sexual dimorphism in defense *δ*_d_. Blue lines indicate the fit of a linear regression model generated using the *lmplot* function from the *seaborn* package in Python 3.12.

## 4 Discussion

We showed that sexual selection can drive the evolution of aposematism, even when prey do not initially have a defense mechanism. Sexual selection can favor the evolution of a signal, which increases prey detectability. This, in turn, increases predation pressure, favoring the evolution of a defense mechanism. When prey display a signal and are defended, predators may then learn to associate the signal with defense and avoid attacking these prey, making the prey signal a ‘warning’ signal. This evolutionary process at the origin of aposematism can be called the ‘sexual selection hypothesis’ for the origin of aposematism.

In addition to demonstrating that sexual selection can lead to the evolution of aposematism, our study highlights key features that can arise when sexual selection is at play. This could help identify empirical cases of aposematism in nature that support the ‘sexual selection hypothesis’ for the origin of aposematism. We show that under sexual selection, the evolution of the signal can precede the evolution of defense (but note that sexual selection can also favor the evolution of aposematism in a defended prey population). It may be possible to investigate whether this has occurred in some empirical cases by reconstructing the ancestral state (*e.g*., Wang and Shaffer, 2008; Forthman and Weirauch, 2018). The evolution of the signal before that of the defense may be peculiar to cases of exaptation, where the aposematic signal has evolved under selective pressures unrelated to aposematism. The involvement of sexual selection and the initial evolution of aposematic signals as sexual signals would constitute a likely evolutionary path to aposematism in these empirical cases. It should be noted that other evolutionary pathways implying exaptation are possible. In particular, a warning signal may also evolve initially as a signal indicating social dominance, thus preventing costly conflicts. Indeed, some badges of status can be colorful (e.g., Pryke et al., 2002; Murphy et al., 2009), although dominance is usually indicated by the color black (Kenyon and Martin, 2023).

Another key feature that arises under the ‘sexual selection hypothesis’ for the origin of aposematism is high sexual dimorphism in signaling and defense. We showed that when sexual selection acts mostly in males, males have not only a stronger signal than females at evolutionary equilibrium, but also a stronger defense. The opposite outcome is expected to occur: when sexual selection acts mostly in females, females may have a stronger signal and defense than males at evolutionary equilibrium. Empirical evidence shows that there is sexual dimorphism in signaling in some aposematic populations (Klein and de Araújo, 2013; Rojas and Endler, 2013; Maan and Cummings, 2009; Palacios-Rodríguez et al., 2022), which could indicate that aposematism was favored by sexual selection in these empirical cases. In contrast, when there is no sexual dimorphism in aposematic signal (e.g., Betancourth-Cundar and Palacios-Rodriguez, 2022), aposematism may have been favored by processes not involving sexual selection. In our model, perhaps surprisingly, sexual dimorphism in defense is extremely strong at evolutionary equilibrium, as females have no defense at all, unlike males. This occurs because the evolution of male signal through sexual selection increases the predation rate (favoring the evolution of male defense), while the evolution of female signal through protection provided by aposematism reduces the predation rate (disfavoring the evolution of female defense). Empirical evidence for sexual dimorphism in defense in aposematic species is lacking, as many studies on variation in defense level do not account for the sex of prey (e.g., Chouteau et al. (2019); Amézquita et al. (2017)). Yet, Arias et al. (2016a) showed in Heliconius butterflies that taking the sex of the prey into account makes it possible to explain a greater variation in prey consumption and therefore in the prey’s level of defense (with, however, greater defense in females than in males, perhaps due to ecological differences between males and females; Arias et al. (2016b)). Our model suggests that assessing the level of sexual dimorphism in defense in aposematic species should be a fruitful line of research, as it could shed light on the initial evolution of aposematism. It is important to note that the observation of sexual dimorphism is not a reliable proof supporting the ‘sexual selection hypothesis’ for the origin of aposematism. Aposematism may first evolve through processes not involving sexual selection (*e.g*., via kin selection in structured populations) and the aposematic signal may have been later used as a sexual signal, leading to sexual dimorphism. Likewise, an aposematic signal may have evolved initially as a sexual signal, but sexual dimorphism may have been lost later as sexual selection was no longer at play.

In our model with sex-dependent traits, we made several simplifying assumptions that favor sexual dimorphism. We expect sexual dimorphism to be less pronounced when these assumptions are not met. We assume a simple genetic architecture where male and female traits have a totally independent genetic basis, whereas this is not necessarily the case in nature. We also assume that only females express mating preferences, yet many studies report male mate choice in aposematic species (Finkbeiner et al., 2014a; Hausmann et al., 2021; Kuo et al., 2024, e.g.,). Finally, we have not taken into account the fact that selective pressures other than sexual selection can act differently on male and female traits, and can determine the intensity of sexual dimorphism. For instance, females and males may have different escape abilities, impacting their predation rate. If females are more predated than males (as shown in a butterfly species; Ohsaki, 1995), this should increase the advantage of females to express the aposematic signal, reducing the intensity of sexual dimorphism in signaling. Similarly, defense and signaling may not be as costly for males as for females (e.g., Blount et al., 2023), which should impact sexual dimorphism. Finally, other selective forces independent of sex (*e.g*. kin selection) could promote the initial evolution of aposematism at the same time as sexual selection, thus reducing the intensity of sexual dimorphism.

In our models, we have assumed that all females express the same mating preference for signaling males. Yet, it has been shown that mating preferences can be genetically encoded and subject to evolution (e.g., in *Heliconius* butterflies; Rossi et al., 2024). If mating preferences evolve, the strength of the resulting sexual selection will determine the initial evolution of signaling, as we have shown that it must outweigh the intrinsic costs for signaling to be favored. Once signaling and defense have both evolved and resulted in aposematism, we do not expect sexual selection to vanish because the evolutionary forces affecting mating preferences should be maintained. For instance, if mating preference is favored as a result of signaling indicating a high viability (under the ‘good-gene hypothesis’), this should still be the case once aposematism has evolved. Whether this occurs for all types of mating preference that lead to sexual selection should be further investigated using theoretical models.

Overall, our theoretical models add support to the idea that sexual selection can play a role in the initial evolution of aposematic signals. We have shown that sexual selection due to mating preferences can favor the initial evolution of signaling, which in turn increases the predation rate and favors the evolution of defense. This exaptation scenario, characterized by a change in the ‘function’ of the signaling trait during evolution, constitutes a likely evolutionary pathway for the evolution of some aposematic signals. Some indirect empirical evidence is likely to confirm the ‘sexual selection hypothesis’ for the origin of aposematism in some aposematic species. Indeed, we have shown that the evolution of signaling before defense, and a high level of sexual dimorphism are expected when sexual selection comes into play in the evolution of aposematism. More generally, our study should stimulate further research on other evolutionary hypotheses for the origin of aposematism involving exaptation.

## 5 Data Availability

Codes are available online at zenodo.org/records/14826541 (DOI 10.5281/zenodo.14826540.).

## Appendix to

### A1 Predator avoidance learning

This section presents how we model the learning dynamics of predators, following Ferreira and Marcon (2014). Initially, predators are naive and attack any prey they encounter. As explained in the main text, predators encounter prey at a rate of *p*. When a predator encounters a prey, it detects the prey with a probability *P*_D_(*s*_•_) depending on the prey’s signal intensity *s*_•_ (see Eq. (2)) and notices the signal with probability *P*_N_(*s*_•_) (see Eq. (3))). Whether or not the naive predator has noticed the signal, it attacks the prey. After attacking a prey whose signal the predator has noticed, the predator may learn to avoid attacking signaling prey with probability *P*_L_(*d*_•_), depending on the prey’s defense level *d*_•_. In line with empirical evidence, we assume that the avoidance learning probability *P*_L_(*d*_•_) increases with the prey’s defense level *d*_•_ (*e.g*. Lindström et al. (1997); Skelhorn and Rowe (2006)), so that

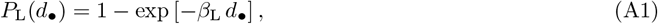

where *β*_L_ modulates the effect of the prey’s defense level *d*_•_ on avoidance learning. We assume that once predators have learned to avoid attacking signaling prey, they will not attack prey when they notice prey’s signal.

We consider a large population of predators and denote *θ* the proportion of experienced predators that have learned to avoid attacking signaling prey. We assume that predators forget at a rate of *λ*. The dynamics of *θ* through time is given by the following differential equation:

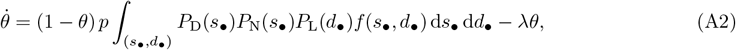

where *f* is the distribution of traits in the prey population.

We assume that the prey population is much larger than the predator population. As a result, the proportion *θ* of experienced predators reaches its equilibrium value 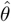 before predation significantly alters the distribution of traits in prey. To investigate the evolutionary dynamics in prey, we then consider the proportion of experienced predators at equilibrium (see Eq. (4)). This equilibrium value is given by

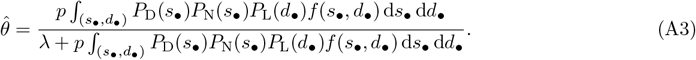

### A2 Individual based simulation

Our individual-based simulations track the evolution of the trait distribution for the model with sex-dependent traits across a fixed number of generations, following the life cycle described in the main text. Each individual *i* at each generation is characterized by the vector of its four traits (*s*_f, *i*_, *d*_f, *i*_, *s*_m,*i*_, *d*_m,*i*_), as well as its sex *χ*_*i*_ and expressed phenotype vector (*s*_*i*_, *d*_*i*_). The expressed phenotype vector of each individual *i* depends on their trait vector and sex, such that (*s*_*i*_, *d*_*i*_) = (*s*_f, *i*_, *d*_f, *i*_) when *χ*_*i*_ = female or (*s*_*i*_, *d*_*i*_) = (*s*_m,*i*_, *d*_m,*i*_) when *χ*_*i*_ = male. To optimize computation time, in the first generation, the values of *s*_f, *i*_, *d*_f, *i*_, *s*_m,*i*_, *d*_m,*i*_ for each individual *i* ∈ {1, …, *N*} are initialized to their values at the evolutionary equilibrium determined from the invasion analyses. Additionally, the sex *χ*_*i*_ of each individual *i* is randomly assigned, with an equal probability of being female or male.

At each generation *g* ≥ 1, the following occurs:

#### (i) Survival

We simulate a sequence of death events using recursion, which modifies the distribution of traits in the population and the population size. Let *t* be the time of the most recent death event, and *N* (*t*) denote the number of adults remaining after that event. Initially, before any death events have occurred, we set *t* = 0 and *N* (0) = *N*. The recursion is then simulated iteratively until *t* ≥ 1

a. We first compute the proportion of experimented predators as (obtained by substituting 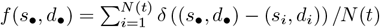, where *δ* is the Dirac function, into Eq. (A3))

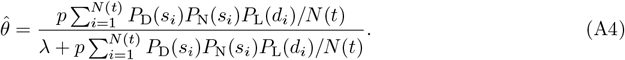
b. For each individual we compute its mortality rate *μ*_*i*_ = *μ*_c_(*s*_*i*_, *d*_*i*_) + *μ*_p_(*s*_*i*_, *d*_*i*_) where the expression of *μ*_c_(*s*_*i*_, *d*_*i*_) and *μ*_p_(*s*_*i*_, *d*_*i*_) are given by eqs. (1) and (8) with (*s*_•_, *d*_•_) = (*s*_*i*_, *d*_*i*_). Note that the predation-induced mortality rate *μ*_*i*_ depends on the proportion of experienced predators defined in Eq. (A4).
c. We simulate an exponential random variable *τ* with parameter 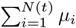. We then update *t* = *t* + *τ*, representing the time of the next event. If *t* ≤ 1, an individual is randomly selected to die, where the probability to select an individual is proportional to their mortality rate *μ*_*i*_. Note that this process is equivalent to associating an independent exponential clock with parameter *μ*_*i*_ to each individual *i*. The first clock to ring determines the individual that dies. The random variable *τ* follows the same probability density as the time at which the first exponential clock rings, and the probability of selecting an individual is equivalent to the probability that their exponential clock rings first. After a death event, the population size is then reduced by 1, and the surviving individuals are renumbered accordingly.

#### (ii) Reproduction

For each male *i*, we compute its attractiveness *A*_*i*_ = 𝒜 (*s*_*i*_) where the expression for 𝒜 (*s*_*i*_) is given by Eq. (14) with *s*_•_ = *s*_*i*_ and 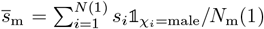. Here *N* (1) and *N*_m_(1) are the number of adults and males alive at the end of the survival stage. Each female selects a male to form a pair, with the probability of choosing a particular male *i* being proportional to his attractiveness *A*_*i*_. Note that a single male may be chosen by multiple females, resulting in his participation in multiple pairs. Next, *N* offspring are produced, with each offspring randomly assigned to a mating pair, meaning that multiple offspring may originate from the same pair. This is equivalent to each pair producing *k* offspring, after which *N* offspring are uniformly sampled from the total pool of offspring to form the next generation. Each mating pair transmits a vector of the four traits to its offspring, with each trait being inherited from either the mother or the father with equal probability. Each offspring was then assigned a sex with equal probability.

#### (iii) Mutation

With probability 1 − *μ*, an offspring retains the traits transmitted by its parents. Otherwise (with probability *μ*), traits mutate; we model this by adding to each inherited trait value an effect sampled from a normal distribution with mean 0 and variance *σ*^2^. If necessary, the resulting trait values are truncated to remain positive. In all simulations, we set *μ* = 5.10^−4^ and *σ* = 5.10^−4^.

We repeat steps (i)-(iii) for a fixed number of generations (see figure legends for parameter values).

### A3 Proportion of experienced predators in the invasion analyses

In the case of a sexually monomorphic population where all prey have the same signal intensity *s* and defense level *d* so that *f* (*s*_•_, *d*_•_) = *δ* ((*s*_•_, *d*_•_) − (*s, d*)), where *δ* is the Dirac function (which is zero everywhere except at (0,0), where it is infinitely high, and the integral of which over the entire phenotypic space is equal to one), the proportion of experienced predators at equilibrium becomes:

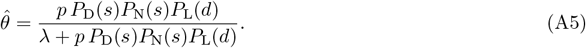

In the case of a sexually dimorphic population where all females (resp. males) have the same signal intensity *s*_f_ (resp. *s*_m_) and defense level *d*_f_ (resp. *d*_m_), and the sex ratio remains constant, the proportion of experienced predators at equilibrium becomes:

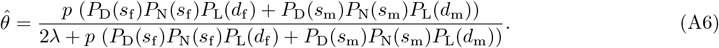

Note that Eq. (A6) relies on the assumption that the distribution of traits remains nearly constant during a generation, resulting in a balanced sex ratio. However, the sex ratio is only balanced at the start of a generation, since males and females experience different death rates due to predation and trait-related costs. We relax this assumption in our individual-based simulations (see Appendix A2 and Fig. A2).

### A4 Selection gradients

#### A4.1 Model with sex-independent traits

From Eq. (23), we determine the selection gradient on each trait. The selection gradient on the signal is given by

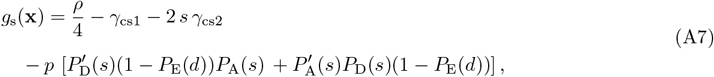

where 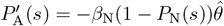 with the expression of 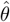 given in Eq. (A5).

The selection gradient on the defense is given by

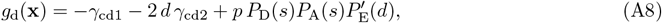

where 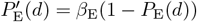.

#### A4.2 Model with sex-dependent traits

From Eq. (23), we determine the selection gradient on each trait, just like with model with sex-independent traits. The selection gradient on the female signal is given by

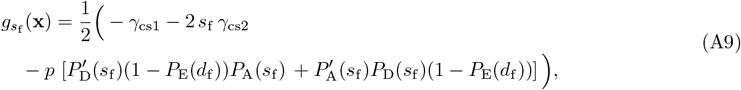

where 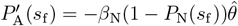 with the expression of 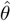 given in Eq. (A6).

The selection gradient on the female defense is given by

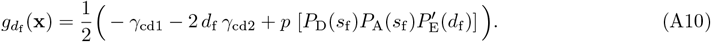

The selection gradient on the male signal is given by

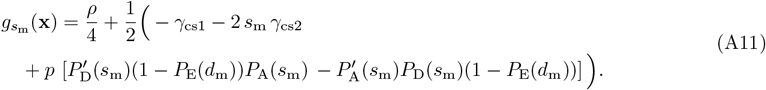

Finally, the selection gradient on the male defense is given by

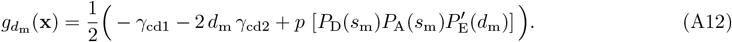

### A5 Sensitivity analyses

In this section, we describe the sensitivity analyses we conduct to evaluate the robustness of our analytical results. We carry out the sensitivity analyses for the models with sex-independent or sex-dependent traits. We begin by randomly selecting the values of the parameters (see below for details on the range of parameter values). For each combination of parameter values drawn at random, we numerically determine the corresponding evolutionary equilibrium. We assume that the population initially displays no signal and no defense (*s* = *d* = 0 or *s*_f_ = *d*_f_ = *s*_m_ = *d*_m_ = 0, depending on the model). We then use Eq. (22) with *Nμσ*^2^ = 0.001 to model the evolution of mean trait values through generations. During the simulations, we constrain the mean trait values to ensure that they remain within the phenotypic space. We run the simulations until the norm of the changes in mean trait values falls below a threshold of 10^−6^. This condition ensures that the simulations have reached a state close to equilibrium, where subsequent changes in mean trait values are negligible.

As mentioned above, all parameter values are randomly drawn within a given range. The parameter *P*_D0_ is randomly drawn from a uniform distribution within the range [0, 1]. For the remaining parameters (*γ*_cs1_, *γ*_cs2_, *γ*_cd1_, *γ*_cd2_, *β*_D_, *β*_N_, *β*_L_, *β*_E_, *λ, ρ, p*), we generate random values by drawing *u* from a uniform distribution within [−2, 2] and then computing 10^*u*^. The use of exponential transformations ensures that parameter values span several orders of magnitude, enabling in-depth exploration of the parameter space.

To analyze the simulated data, we compute the Pearson correlation coefficient between parameters and variables (see Tables A1 and A2). We compute the mean absolute correlation between parameter values in the dataset comprising 1,000,000 simulation outputs. If there was an infinite number of replicates, this correlation should be zero because the parameters are drawn independently of each other. Any correlation between parameters and variables whose absolute value is not one order of magnitude greater than the mean absolute correlation between parameters is considered not to be significant. Indeed, we cannot distinguish whether the correlation is due to the parameters affecting the variables at evolutionary equilibrium or to correlations between parameters in the dataset.

### A6 Selection on the signal in the ancestral population

We assume that the ancestral population has reached a state of evolutionary equilibrium, with sexual selection playing no role (*ρ* = 0). Here, we show that in the ancestral population the signal is always counter-selected.

The selection gradient on the signal describes whether a higher signal intensity is favored by natural selection. In the model with sex-independent traits, the selection gradient on the signal is initially equal to

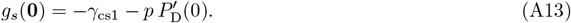

This is obtained by substituting *s* = *d* = 0 and *ρ* = 0 into Eq. (A7). Since *g*_*s*_(**0**) is always negative, the signal is always counter-selected.

In the model with sex-dependent traits, the selection gradients on female and male signals are initially equal to

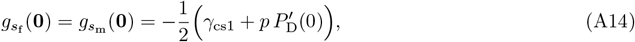

This is obtained by substituting *s*_f_ = *d*_f_ = *s*_m_ = *d*_m_ = 0 and *ρ* = 0 into Eqs. (A9) and (A11). Since 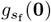 and 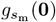 are always negative, the male and female signals are always counter-selected.

### A7 Initial evolution in the model with sex-independent traits

#### A7.1 Initial evolution of the signal

Here, we derive Eq. (24) of the main text. This gives the condition for the initial evolution of the signal in a population characterized by *s* = 0 and *d* = 0 in the model with sex-independent traits. The signal can initially evolve if

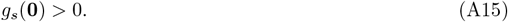

By substituting the expression of *g*_*s*_(**0**) (obtained by substituting *s* = *d* = 0 into Eq. (A7)) into the condition (A15), we get

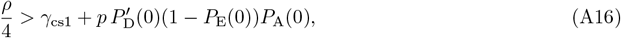

which is Eq. (24) in the main text.

#### A7.2 Initial evolution of the defense

Here, we derive Eq. (25) of the main text. This gives the condition for the initial evolution of the defense in a population characterized by *s >* 0 and *d* = 0 in the model with sex-independent traits. The defense can initially evolve if

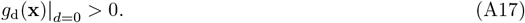

By substituting the expression of *g*_d_(**x**)|_*d*=0_ (obtained by substituting *d* = 0 into Eq. (A8)) into the condition (A17), we get

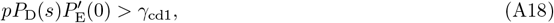

which is Eq. (24) in the main text.

From Eq. (A18), we derive Eq. (26) of the main text, which gives the threshold value 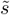 that signal intensity must exceed for predation pressure to be high enough to favor defense evolution. By substituting the expression of *P*_D_(*s*) (obtained by substituting *s*_•_ = *s* into Eq. (2)) into Eq. (A18) and isolating *s*, we get the condition

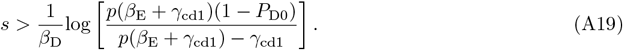

The threshold value is therefore

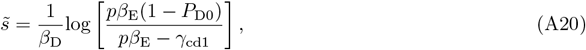

which is Eq. (26) in the main text.

### A8 Initial evolution in the model with sex-dependent traits

#### A8.1 Initial evolution of male signal and defense

Here, we derive the condition for the initial evolution of male signal and defense in the model with sex-dependent traits. In a population characterized by *s*_f_ = 0, *d*_f_ = 0, *s*_m_ = 0 and *d*_m_ = 0, the male signal can initially evolve if

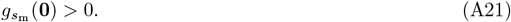

By substituting the expression of 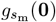 (obtained by substituting *s*_f_ = *d*_f_ = *s*_m_ = *d*_m_ = 0 into Eq. (A11)) into the condition (A21), we get

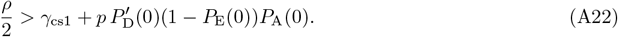

After the initial evolution of male signal (*i.e*., in a population characterized by *s*_f_ = 0, *d*_f_ = 0, *s*_m_ *>* 0 and *d*_m_ = 0) the male defense can initially evolve if

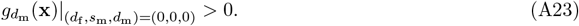

By substituting the expression of 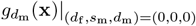 (obtained by substituting *d*_f_ = 0, *s*_m_ = 0 and *d*_m_ = 0 into Eq. (A12)) into the condition (A23), we get

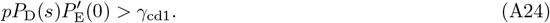

#### A8.2 Initial evolution of the female signal

Here, we derive Eq. (24) of the main text, which gives the condition for the initial evolution of female signal in a population characterized by *s*_f_ = 0, *d*_f_ = 0, *s*_m_ *>* 0 and *d*_m_ *>* 0. The female signal can initially evolve if

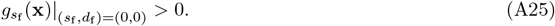

By substituting the expression of 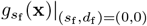 (obtained by substituting *s*_f_ = *d*_f_ = 0 into Eq. (A9)) into the condition (A25), we get

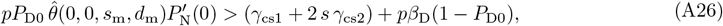

which is Eq. (29) in the main text.

#### A8.3 No evolution of the female defense

Here, we prove that female defense can never evolve in an ancestrally defenseless population, even when the other traits have evolved. Once male signal, male defense and female signal have evolved (*i.e*., in a population characterized by *s*_f_ *>* 0, *d*_f_ = 0, *s*_m_ *>* 0 and *d*_m_ *>* 0; note that *s*_f_ is not necessarily at equilibrium), the female defense cannot evolve if

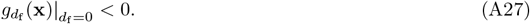

To demonstrate that the condition in Eq. (A27) always holds, we first need to establish that *P*_A_(*s*_f_)*P*_D_(*s*_f_) *< P*_A_(0)*P*_D_(0) = *P*_D0_. This is due to the impact of the female signal on the predation pressure on females. We assume that the male traits reached their evolutionary equilibrium before female signal began to evolve. Throughout the evolution of a more intense female signal, natural selection consistently favors an increase in female signal intensity. In other words, we have

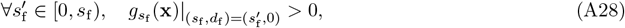

where 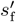 takes all the past values of the variable *s*_f_.

By substituting the expression of the selection gradient on female signal (obtained by substituting 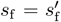 and *d*_f_ = 0 into Eq (A9)), this is equivalent to the condition:

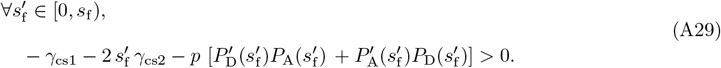

This entails that throughout the evolution of a more intense female signal, we have

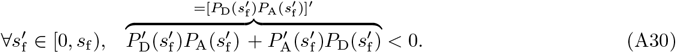

This means that the product 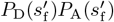 decreases as the female signal evolves, which implies that *P*_A_(*s*_f_)*P*_D_(*s*_f_) *< P*_A_(0)*P*_D_(0), with *P*_A_(0)*P*_D_(0) = *P*_D0_.

Using *P*_A_(*s*_f_)*P*_D_(*s*_f_) *< P*_D0_ and the expression for 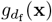 from Eq. (A10), we can show that

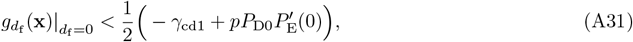

and therefore

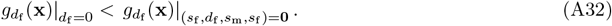

If the population is ancestrally defenseless, we have 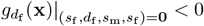, which implies that

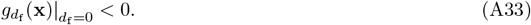

This demonstrates that female defense can never evolve in an ancestrally defenseless population, even when the other traits have evolved.

### A9 Expressions of the levels of sexual dimorphism

In the model with sex-dependent traits, we express the level of sexual dimorphism in signal as

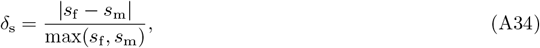

and the level of sexual dimorphism in defense as

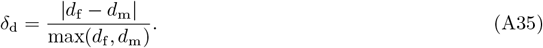

## A10 Supplementary figures and tables

**Figure A1:**
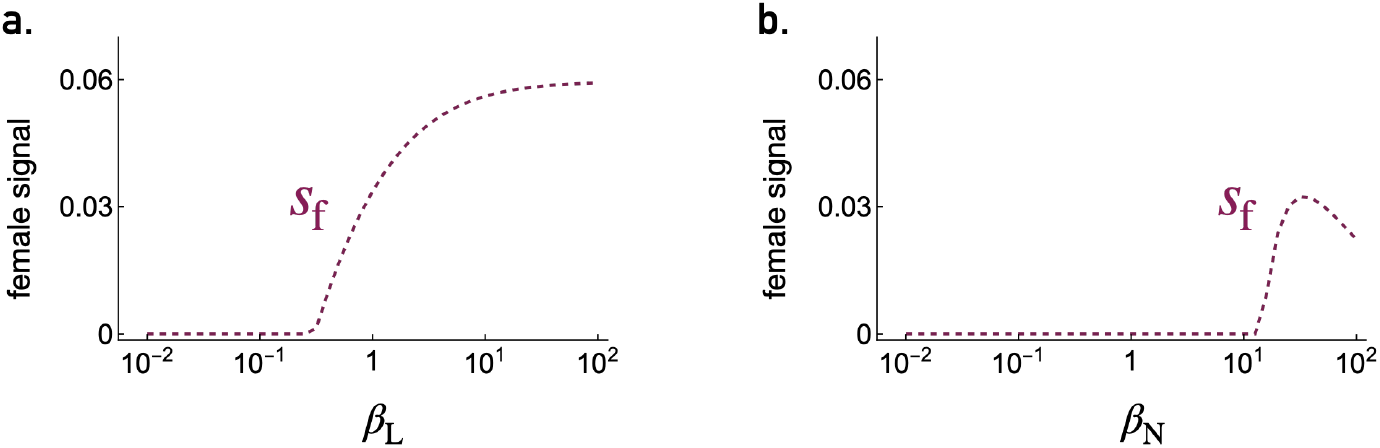
Factors driving the evolution of female signal. Impact on female signal of **a**. the extent to which the defense level increases predator avoidance learning, and **b**. the extent to which the signal intensity increases the probability of the signal being noticed *β*_N_. Parameter values are: *ρ* = 1.2, *γ*_cs1_ = 0.1, *γ*_cs2_ = 0.35, *γ*_cd1_ = 0.08, *γ*_cd2_ = 0.1, *β*_D_ = 1.25, *β*_N_ = 17.5, *β*_E_ = 0.9, *β*_L_ = 0.5, *P*_D0_ = 0.25, *λ* = 0.01, *p* = 0.3.

**Figure A2:**
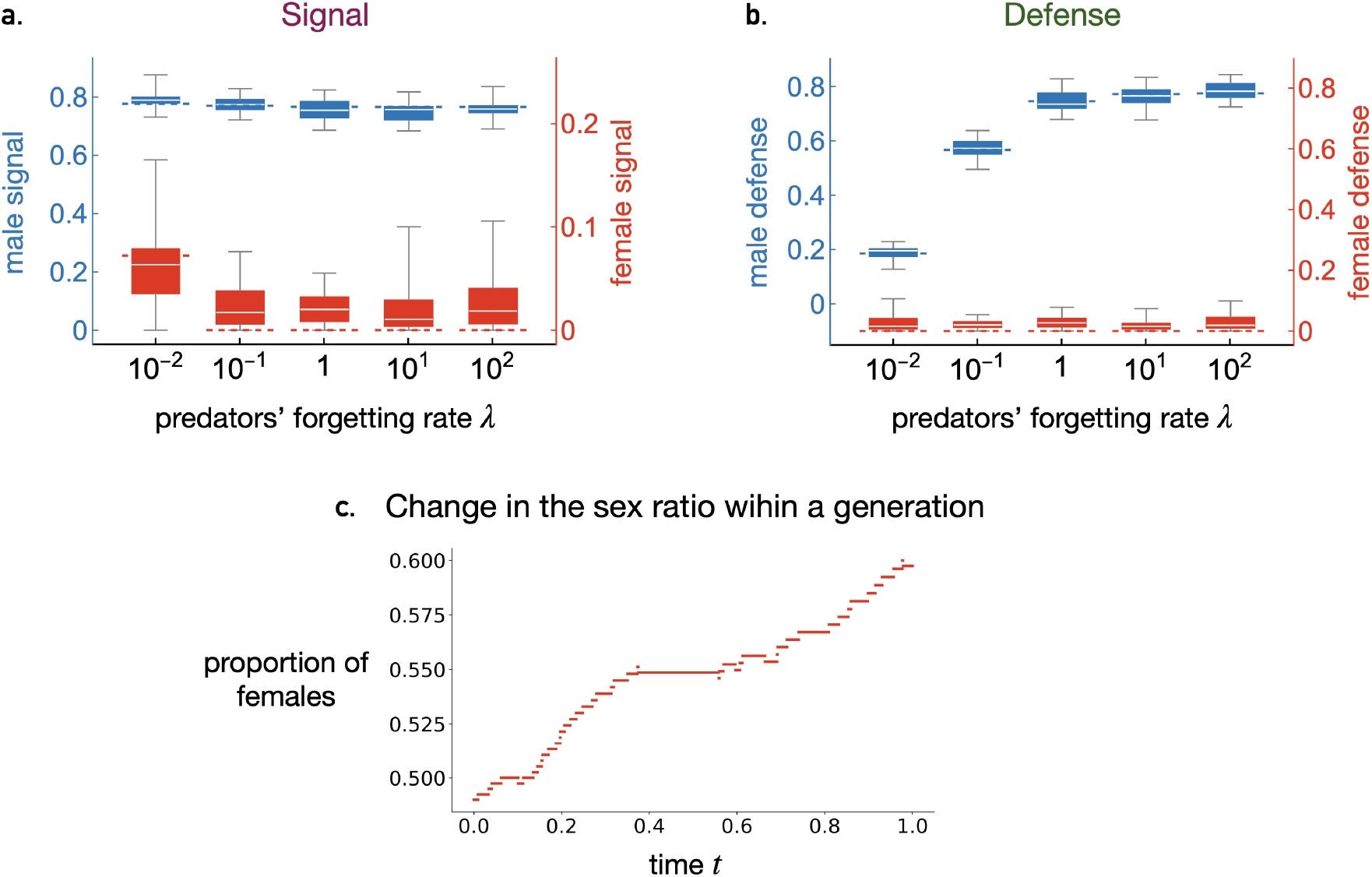
Evolution of male and female signal and defense from individual-based simulations. Box plots show distribution of **a**. signal and **b**. defense over 100,000 generations of 10 replicates at equilibrium from individual-based simulations for different values of the predators’ forgetting rate *λ*. For each *λ*, the population was first allowed to evolve for 150,000 generations to ensure equilibrium was reached. Dashed lines show the evolutionary equilibrium determined from the invasion analyses. **c**. Proportion of females over time within one generation at equilibrium, based on individual-based simulations (*λ* = 1). Parameter values are: *N* = 200, *ρ* = 1.2, *γ*_cs1_ = 0.1, *γ*_cs2_ = 0.35, *γ*_cd1_ = 0.08, *γ*_cd2_ = 0.1, *β*_D_ = 1.25, *β*_N_ = 17.5, *β*_E_ = 0.9, *β*_L_ = 0.5, *P*_D0_ = 0.25, *p* = 0.3.

**Table A1:** Results of the sensitivity analyses performed using the model with sex-independent traits. Pearson correlation coefficient between the different parameters and variables using the dataset produced as described in the section A5 with 1,000,000 combinations of randomly selected parameter values. The mean absolute correlation between parameters is equal to 0.00077 ≃ 0.001. The symbol ^∗^ indicates the correlation coefficients whose absolute values are an order of magnitude greater than this mean absolute correlation between parameters, and which we therefore consider significant. This implies that we assume these parameters are likely to affect these variables at evolutionary equilibrium. For the other correlations (without the symbol ^∗^), we cannot distinguish whether the correlation is due to the parameters affecting the variables at evolutionary equilibrium, or due to the correlations between the parameters in the dataset.

| | $s$ | $d$ | $\hat{\theta}$ | $1 - \mathcal{P}(s, d)$ |
| --- | --- | --- | --- | --- |
| $\gamma_{cs1}$ | -0.07697* | 0.00695 | -0.10196* | -0.00903 |
| $\gamma_{cs2}$ | -0.08493* | 0.00704 | -0.04105* | -0.00247 |
| $\gamma_{cd1}$ | -0.00137 | -0.10873* | -0.08633* | 0.12014* |
| $\gamma_{cd2}$ | -0.00134 | -0.10331* | -0.02571* | 0.06935* |
| $\beta_D$ | -0.02529* | 0.00619 | -0.06544* | -0.00282 |
| $\beta_N$ | 0.00354 | -0.02815* | 0.13474* | -0.00493 |
| $\beta_L$ | -0.00037 | -0.00517 | 0.06584* | -0.00540 |
| $\beta_E$ | 0.00609 | -0.07704* | 0.03362* | -0.30648* |
| $\lambda$ | 0.00042 | 0.01269* | -0.10708* | 0.00500 |
| $\rho$ | 0.34907* | -0.00503 | 0.11881* | 0.02098* |
| $P_{D0}$ | 0.01195* | 0.08132* | 0.10365* | 0.15876* |
| $p$ | -0.01375* | 0.29290* | 0.29116* | 0.48017* |

**Table A2:** Results of the sensitivity analyses performed using the model with sex-dependent traits. Pearson correlation coefficient between the different parameters and variables using the dataset produced as described in the section A5 with 1,000,000 combinations of randomly selected parameter values. The mean absolute correlation between parameters is equal to 0.0025. The symbol ^∗^ indicates the correlation coefficients whose absolute values are an order of magnitude greater than this mean absolute correlation between parameters, and which we therefore consider significant. This implies that we assume these parameters are likely to affect these variables at evolutionary equilibrium. For the other correlations (without the symbol ^∗^), we cannot distinguish whether the correlation is due to the parameters affecting the variables at evolutionary equilibrium, or due to the correlations between the parameters in the dataset.

| | $s_f$ | $d_f$ | $s_m$ | $d_m$ | $\delta_s$ | $\delta_d$ | $\hat{\theta}$ | $1 - \mathcal{P}(s_m, d_m)$ | $1 - \mathcal{P}(s_f, d_f)$ |
| --- | --- | --- | --- | --- | --- | --- | --- | --- | --- |
| $\gamma_{cs1}$ | -0.03020* | 0.00006 | -0.07442* | -0.00059 | -0.29974* | -0.10513* | -0.09657* | -0.01281* | -0.00147 |
| $\gamma_{cs2}$ | -0.02706* | -0.00025 | -0.09003* | -0.00089 | -0.00743 | -0.06784* | -0.04529* | -0.00813 | 0.00065 |
| $\gamma_{cd1}$ | -0.01094* | -0.10807* | -0.00161 | -0.10906* | -0.01744* | -0.08227* | -0.07187* | 0.12557* | 0.12102* |
| $\gamma_{cd2}$ | 0.00110 | -0.10145* | 0.00467 | -0.10229* | -0.01093* | 0.00090 | -0.03221* | 0.07211* | 0.06993* |
| $\beta_D$ | -0.02768* | 0.00271 | -0.02877* | 0.00210 | -0.14745* | -0.04743* | -0.05264* | -0.00393 | 0.00300 |
| $\beta_N$ | -0.01317* | -0.00628 | 0.00061 | -0.00461 | -0.00186 | 0.00487 | 0.03285* | -0.00281 | -0.00445 |
| $\beta_L$ | 0.01173* | 0.00353 | 0.00232 | 0.00223 | -0.00222 | 0.01420* | 0.05852* | -0.00056 | 0.00072 |
| $\beta_E$ | -0.01479* | -0.07955* | 0.01330* | -0.07980* | 0.06580* | 0.06258* | 0.08187* | -0.30477* | -0.29621* |
| $\lambda$ | -0.02384* | 0.00346 | 0.00231 | 0.00789 | 0.00686 | -0.01004* | -0.09200* | 0.00610 | 0.00349 |
| $\rho$ | 0.03299* | -0.00427 | 0.37083* | -0.00314 | 0.46450* | 0.27812* | 0.19809* | 0.02848* | -0.00286 |
| $P_{D0}$ | 0.04383* | 0.08725* | 0.00402 | 0.08012* | 0.05941* | -0.07959* | 0.05990* | 0.15079* | 0.17489* |
| $p$ | 0.08723* | 0.30479* | -0.01784* | 0.30191* | -0.11879* | 0.00680 | 0.18184* | 0.48866* | 0.50206* |

## References

Amézquita, A., Ramos, O., González, M. C., Rodríguez, C., Medina, I., Simoes, P. I., and Lima, A. P. (2017). Conspicuousness, color resemblance, and toxicity in geographically diverging mimicry: The pan-amazonian frog allobates femoralis. Evolution, 71(4):1039–1050.

Arias, M., Mappes, J., Théry, M., and Llaurens, V. (2016a). Inter-species variation in unpalatability does not explain polymorphism in a mimetic species. Evolutionary Ecology, 30(3):419–433.

Arias, M., Meichanetzoglou, A., Elias, M., Rosser, N., de Silva, D. L., Nay, B., and Llaurens, V. (2016b). Variation in cyanogenic compounds concentration within a heliconius butterfly community: does mimicry explain everything? BMC Evolutionary Biology, 16(1):272.

Aubier, T. G. and Sherratt, T. N. (2015). Diversity in müllerian mimicry: The optimal predator sampling strategy explains both local and regional polymorphism in prey. Evolution, 69(11):2831–2845.

Benson, W. W. (1971). Evidence for the evolution of unpalatability through kin selection in the heliconinae (lepidoptera). The American Naturalist, 105(943):213–226.

Betancourth-Cundar, M. and Palacios-Rodriguez, P. (2022). Reproductive behaviors promote ecological and phenotypic sexual differentiation in the critically endangered lehmann’s poison frog. Evolutionary Ecology, 36(6):1077–1093.

Blount, J. D., Rowland, H. M., Drijfhout, F. P., Endler, J. A., Inger, R., Sloggett, J. J., Hurst, G. D. D., Hodgson, D. J., and Speed, M. P. (2012). How the ladybird got its spots: effects of resource limitation on the honesty of aposematic signals. Functional Ecology, 26(2):334–342.

Blount, J. D., Rowland, H. M., Mitchell, C., Speed, M. P., Ruxton, G. D., Endler, J. A., and Brower, L. P. (2023). The price of defence: toxins, visual signals and oxidative state in an aposematic butterfly. Proceedings of the Royal Society B: Biological Sciences, 290(1991):20222068.

Boussens-Dumon, G. and Llaurens, V. (2021). Sex, competition and mimicry: an eco-evolutionary model reveals unexpected impacts of ecological interactions on the evolution of phenotypes in sympatry. Oikos, 130(11):2028–2039.

Broom, M., Ruxton, G. D., and Speed, M. P. (2008). Evolutionarily Stable Investment in Anti-Predatory Defences and Aposematic Signalling, pages 37–48. Birkhäuser Boston, Boston, MA.

Broom, M., Speed, M., and Ruxton, G. (2006). Evolutionarily stable defence and signalling of that defence. Journal of Theoretical Biology, 242(1):32–43.

Bryant, J. and Julkunentiitto, R. (1995). Ontogenic development of chemical defense by seedling resin birch - energy-cost of defense production. Journal of chemical ecology, 21(7):883–896.

Champagnat, N., Ferrière, R., and Méléard, S. (2006). Unifying evolutionary dynamics: From individual stochastic processes to macroscopic models. Theoretical Population Biology, 69(3):297–321. ESS Theory Now.

Chouteau, M., Dezeure, J., Sherratt, T. N., Llaurens, V., and Joron, M. (2019). Similar predator aversion for natural prey with diverse toxicity levels. Animal Behaviour, 153:49–59.

Chouteau, M., Llaurens, V., Piron-Prunier, F., and Joron, M. (2017). Polymorphism at a mimicry supergene maintained by opposing frequency-dependent selection pressures. Proceedings of the National Academy of Sciences of the United States of America, 114(31):8325–8329.

Cortesi, F. and Cheney, K. L. (2010). Conspicuousness is correlated with toxicity in marine opisthobranchs. Journal of Evolutionary Biology, 23(7):1509–1518.

Dale, J., Dey, C. J., Delhey, K., Kempenaers, B., and Valcu, M. (2015). The effects of life history and sexual selection on male and female plumage colouration. Nature, 527(7578):367–370.

Dieckmann, U. and Law, R. (1996). The dynamical theory of coevolution: a derivation from stochastic ecological processes. Journal of mathematical biology, 34:579–612.

Estrada, C. and Jiggins, C. D. (2008). Interspecific sexual attraction because of convergence in warning colouration: is there a conflict between natural and sexual selection in mimetic species? Journal of Evolutionary Biology, 21(3):749–760.

Ferreira, W. C. and Marcon, D. (2014). Revisiting the 1879 model for evolutionary mimicry by fritz müller: New mathematical approaches. Ecological Complexity, 18:25–38. Progress in Mathematical Population Dynamics and Ecology (Proceedings of MPDE 2012).

Finkbeiner, S. D., Briscoe, A. D., and Reed, R. D. (2014a). Warning signals are seductive: Relative contributions of color and pattern to predator avoidance and mate attraction in heliconius butterflies. Evolution, 68(12):3410–3420.

Finkbeiner, S. D., Briscoe, A. D., and Reed, R. D. (2014b). Warning signals are seductive: Relative contributions of color and pattern to predator avoidance and mate attraction in Heliconius butterflies. Evolution, 68(12):3410–3420.

Fisher, R. A. (1930). The genetical theory of natural selection,. The Clarendon Press, Oxford.

Forthman, M. and Weirauch, C. (2018). Phylogenetic comparative analysis supports aposematic colouration–body size association in millipede assassins (Hemiptera: Reduviidae: Ectrichodiinae). Journal of Evolutionary Biology, 31(7):1071–1078.

Franks, D. W., Ruxton, G. D., and Sherratt, T. N. (2009). Warning signals evolve to disengage batesian mimics. Evolution, 63(1):256–267.

Gittleman, J. L. and Harvey, P. H. (1980). Why are distasteful prey not cryptic? Nature, 286(5769):149–150.

Gordon, S. P., Kokko, H., Rojas, B., Nokelainen, O., and Mappes, J. (2015). Colour polymorphism torn apart by opposing positive frequency-dependent selection, yet maintained in space. Journal of Animal Ecology, 84(6):1555–1564.

Grill, C. P. (1999). Development of colour in an aposematic ladybird beetle: the role of environmental conditions. Evolutionary Ecology Research, 1(6):651–662.

Guilford, T. (1985). Is kin selection involved in the evolution of warning coloration? Oikos, 45(1):31–36.

Halpin, C. G., Skelhorn, J., and Rowe, C. (2008a). Being conspicuous and defended: selective benefits for the individual. Behavioral Ecology, 19(5):1012–1017.

Halpin, C. G., Skelhorn, J., and Rowe, C. (2008b). Naïve predators and selection for rare conspicuous defended prey: the initial evolution of aposematism revisited. Animal Behaviour, 75(3):771–781.

Harvey, P. H., Bull, J. J., Pemberton, M., and Paxton, R. J. (1982). The evolution of aposematic coloration in distasteful prey: A family model. The American Naturalist, 119(5):710–719.

Hausmann, A. E., Kuo, C.-Y., Freire, M., Rueda-M, N., Linares, M., Pardo-Diaz, C., Salazar, C., and Merrill, R. M. (2021). Light environment influences mating behaviours during the early stages of divergence in tropical butterflies. Proceedings of the Royal Society B: Biological Sciences, 288(1947):20210157.

Hedley, E. and Caro, T. (2022). Aposematism and mimicry in birds. Ibis, 164(2):606–617.

Hill, G. E. (1991). Plumage coloration is a sexually selected indicator of male quality. Nature, 350(6316):337– 339.

Howell, N., Sheard, C., Koneru, M., Brockelsby, K., Ono, K., and Caro, T. (2021). Aposematism in mammals. Evolution, 75(10):2480–2493.

Jiggins, C. D., Naisbit, R. E., Coe, R. L., and Mallet, J. (2001). Reproductive isolation caused by colour pattern mimicry. Nature, 411(6835):302–305.

Järvi, T., Sillén-Tullberg, B., and Wiklund, C. (1981). Individual versus kin selection for aposematic coloration: A reply to harvey and paxton. Oikos, 37(3):393–395.

Kenyon, H. L. and Martin, P. R. (2023). Color as an interspecific badge of status: A comparative test. The American Naturalist.

Klein, A. L. and de Araújo, A. M. (2013). Sexual size dimorphism in the color pattern elements of two mimetic heliconius butterflies. Neotropical Entomology, 42(6):600–606.

Kuo, C.-Y., Melo-Flores, L., Aragon, A., Oberweiser, M. M., McMillan, W. O., Pardo-Diaz, C., Salazar, C., and Merrill, R. M. (2024). Divergent warning patterns influence male and female mating behaviours in a tropical butterfly. Journal of Evolutionary Biology, 37(3):267–273.

Lee, T. J., Marples, N. M., and Speed, M. P. (2010). Can dietary conservatism explain the primary evolution of aposematism? Animal Behaviour, 79(1):63–74.

Leimar, O., Enquist, M., and Sillen-Tullberg, B. (1986). Evolutionary stability of aposematic coloration and prey unprofitability: A theoretical analysis. The American Naturalist, 128(4):469–490.

Lev-Yadun, S. (2009). Müllerian mimicry in aposematic spiny plants. Plant Signaling & Behavior, 4(6):482– 483. PMID: 19816137.

Lev-Yadun, S. (2024). Visual-, olfactory-, and nectar-taste-based flower aposematism. Plants, 13(3).

Lindstedt, C., Boncoraglio, G., Cotter, S., Gilbert, J., and Kilner, R. M. (2017). Aposematism in the burying beetle? dual function of anal fluid in parental care and chemical defense. Behavioral Ecology, 28(6):1414–1422.

Lindstedt, C., Talsma, J. H. R., Ihalainen, E., Lindström, L., and Mappes, J. (2010). Diet quality affects warning coloration indirectly: excretion costs in a generalist herbivore. Evolution, 64(1):68–78.

Lindström, L., Alatalo, R. V., and Mappes, J. (1997). Imperfect batesian mimicry—the effects of the frequency and the distastefulness of the model. Proceedings of the Royal Society of London. Series B: Biological Sciences, 264(1379):149–153.

Maan, M. E. and Cummings, M. E. (2008). Female preferences for aposematic signal components in a polymorphic poison frog. Evolution, 62(9):2334–2345.

Maan, M. E. and Cummings, M. E. (2009). Sexual dimorphism and directional sexual selection on aposematic signals in a poison frog. Proceedings of the National Academy of Sciences, 106(45):19072–19077.

Maan, M. E. and Cummings, M. E. (2012). Poison frog colors are honest signals of toxicity, particularly for bird predators. The American Naturalist, 179(1):E1–E14. PMID: 22173468.

Maan, M. E. and Sefc, K. M. (2013). Colour variation in cichlid fish: developmental mechanisms, selective pressures and evolutionary consequences. In Seminars in Cell & Developmental Biology, volume 24, pages 516–528. Elsevier.

Maisonneuve, L., Chouteau, M., Joron, M., and Llaurens, V. (2021). Evolution and genetic architecture of disassortative mating at a locus under heterozygote advantage. Evolution, 75(1):149–165.

Maisonneuve, L., Elias, M., Smadi, C., and Llaurens, V. (2023). The limits of evolutionary convergence in sympatry: Reproductive interference and historical constraints leading to local diversity in warning traits. The American Naturalist, 201(5):E110–E126.

Mallet, J. and Singer, M. C. (2008). Individual selection, kin selection, and the shifting balance in the evolution of warning colours: the evidence from butterflies. Biological Journal of the Linnean Society, 32(4):337–350.

Marak, H., Biere, A., and Van Damme, J. (2003). Fitness costs of chemical defense in Plantago lanceolata l.:: Effects of nutrient and competition stress. Evolution, 57(11):2519–2530.

Metz, J. a. J. (2011). Thoughts on the geometry of meso-evolution: collecting mathematical elements for a post-modern synthesis. In Chalub, F. A. C. C.and Rodrigues, J., editors, The mathematics of Darwin’s legacy, pages 193–231. Birkhäuser, Basel.

Mullon, C. and Lehmann, L. (2019). An evolutionary quantitative genetics model for phenotypic (co)variances under limited dispersal, with an application to socially synergistic traits. Evolution, 73(9):1695–1728.

Murphy, T. G., Rosenthal, M. F., Montgomerie, R., and Tarvin, K. A. (2009). Female american goldfinches use carotenoid-based bill coloration to signal status. Behavioral Ecology, 20(6):1348–1355.

Nokelainen, O., Hegna, R. H., Reudler, J. H., Lindstedt, C., and Mappes, J. (2012). Trade-off between warning signal efficacy and mating success in the wood tiger moth. Proceedings of the Royal Society B-Biological Sciences, 279(1727):257–265.

Ohsaki, N. (1995). Preferential predation of female butterflies and the evolution of batesian mimicry. Nature, 378(6553):173–175.

Ojala, K., Lindström, L., and Mappes, J. (2007). Life-history constraints and warning signal expression in an arctiid moth. Functional Ecology, 21(6):1162–1167.

Olsson, M., Stuart-Fox, D., and Ballen, C. (2013). Genetics and evolution of colour patterns in reptiles. In Seminars in cell & developmental biology, volume 24, pages 529–541. Elsevier.

Palacios-Rodríguez, P., González-Santoro, M., Amézquita, A., and Brunetti, A. E. (2022). Sexual dichromatism in a cryptic poison frog is correlated with female tadpole transport. Evolutionary Ecology, 36(1):153– 162.

Ponkshe, A. and Endler, J. A. (2022). Joint effects of female preference intensity and frequency-dependent predation on the polymorphism maintenance in aposematic sexual traits. Ecology and Evolution, 12(10).

Pryke, S. R., Andersson, S., Lawes, M. J., and Piper, S. E. (2002). Carotenoid status signaling in captive and wild red-collared widowbirds: independent effects of badge size and color. Behavioral Ecology, 13(5):622– 631.

Reynolds, R. G. and Fitzpatrick, B. M. (2007). Assortative mating in poison-dart frogs based on an ecologically important trait. Evolution, 61(9):2253–2259.

Rigby, M. C. and Jokela, J. (2000). Predator avoidance and immune defence: costs and trade–offs in snails. Proceedings of the Royal Society of London. Series B: Biological Sciences, 267(1439):171–176.

Ripa, J. and Dieckmann, U. (2013). Mutant invasions and adaptive dynamics in variable environments. Evolution, 67(5):1279–1290.

Rojas, B., Burdfield-Steel, E., De Pasqual, C., Gordon, S., Hernandez, L., Mappes, J., Nokelainen, O., Ronka, K., and Lindstedt, C. (2018). Multimodal aposematic signals and their emerging role in mate attraction. Frontiers in Ecology and Evolution, 6.

Rojas, B. and Endler, J. A. (2013). Sexual dimorphism and intra-populational colour pattern variation in the aposematic frog ¡i¿dendrobates tinctorius¡/i¿. Evolutionary Ecology, 27(4, SI):739–753.

Rossi, M., Hausmann, A. E., Alcami, P., Moest, M., Roussou, R., Belleghem, S. M. V., Wright, D. S., Kuo, C.-Y., Lozano-Urrego, D., Maulana, A., Melo-Flórez, L., Rueda-Muñoz, G., McMahon, S., Linares, M., Osman, C., McMillan, W. O., Pardo-Diaz, C., Salazar, C., and Merrill, R. M. (2024). Adaptive introgression of a visual preference gene. Science, 383(6689):1368–1373.

Ruxton, G., Sherratt, T., and Speed, M. (2004). Avoiding Attack: The Evolutionary Ecology of Crypsis, Warning Signals and Mimicry. Oxford biology readers. plOUP Oxford.

Scaramangas, A. and Broom, M. (2022). Aposematic signalling in prey-predator systems: determining evolutionary stability when prey populations consist of a single species. Journal of Mathematical Biology, 85(2):13.

Scaramangas, A., Broom, M., Ruxton, G. D., and Rouviere, A. (2023). Evolutionarily stable levels of aposematic defence in prey populations. Theoretical Population Biology, 153:15–36.

Servedio, M. R. (2000). The effects of predator learning, forgetting, and recognition errors on the evolution of warning coloration. Evolution, 54(3):751–763.

Sherratt, T. N. and Franks, D. W. (2005). Do unprofitable prey evolve traits that profitable prey find difficult to exploit? Proceedings of the Royal Society B: Biological Sciences, 272(1579):2441–2447.

Sillén-Tullberg, B. (1985). Higher survival of an aposematic than of a cryptic form of a distasteful bug. Oecologia, 67(3):411–415.

Sillén-Tullberg, B. and Bryant, E. H. (1983). The evolution of aposematic coloration in distasteful prey: An individual selection model. Evolution, 37(5):993–1000.

Skelhorn, J. and Rowe, C. (2006a). Avian predators taste–reject aposematic prey on the basis of their chemical defence. Biology Letters, 2(3):348–350.

Skelhorn, J. and Rowe, C. (2006b). Prey palatability influences predator learning and memory. Animal Behaviour, 71(5):1111–1118.

Skelhorn, J. and Rowe, C. (2006c). Taste-rejection by predators and the evolution of unpalatability in prey. Behavioral Ecology and Sociobiology, 60(4):550–555.

Stankowich, T., Caro, T., and Cox, M. (2011). Bold coloration and the evolution of aposematism in terrestrial carnivores. Evolution, 65(11):3090–3099.

Stuart–Fox, D. M. and Ord, T. J. (2004). Sexual selection, natural selection and the evolution of dimorphic coloration and ornamentation in agamid lizards. Proceedings of the Royal Society of London. Series B: Biological Sciences, 271(1554):2249–2255.

Tazzyman, S. J. and Iwasa, Y. (2010). Sexual selection can increase the effect of random genetic drift-a quantitative genetic model of polymorphism in Oophaga pumilio, the strawberry poison-dart frog. Evolution, 64(6):1719–1728.

Toledo, L., Sazima, I., and Haddad, C. (2011). Behavioural defences of anurans: an overview. Ethology Ecology & Evolution, 23(1):1–25.

Tullrot, A. (1994). The evolution of unpalatability and warning coloration in soft-bodied marine invertebrates. Evolution, 48(3):925–928.

Waldman, B. and Adler, K. (1979). Toad tadpoles associate preferentially with siblings. Nature, 282(5739):611–613.

Wang, I. J. and Shaffer, H. B. (2008). Rapid color evolution in an aposematic species: A phylogenetic analysis of color variation in the strikingly polymorphic strawberry poison-dart frog. Evolution, 62(11):2742–2759.

Wiklund, C. and Järvi, T. (1982). Survival of distasteful insects after being attacked by naive birds: A reappraisal of the theory of aposematic coloration evolving through individual selection. Evolution, 36(5):998– 1002.

Yeager, J. and Penacchio, O. (2023). Outcomes of multifarious selection on the evolution of visual signals. Proceedings of the Royal Society B: Biological Sciences, 290(1996):20230327.

Zalucki, M. P., Malcolm, S. B., Paine, T. D., Hanlon, C. C., Brower, L. P., and Clarke, A. R. (2001). It’s the first bites that count: Survival of first-instar monarchs on milkweeds. Austral Ecology, 26(5):547–555.

## References

Skelhorn, J. and Rowe, C. (2006). Prey palatability influences predator learning and memory. Animal Behaviour, 71(5):1111–1118.

